# Genetic and developmental divergence in the neural crest programme between cichlid fish species

**DOI:** 10.1101/2024.01.30.578004

**Authors:** Aleksandra Marconi, Grégoire Vernaz, Achira Karunaratna, Maxon J. Ngochera, Richard Durbin, M. Emília Santos

## Abstract

Neural crest (NC) is a vertebrate-specific embryonic progenitor cell population at the basis of important vertebrate features such as the craniofacial skeleton and pigmentation patterns. Despite the wide-ranging variation of NC-derived traits across vertebrates, the contribution of NC to species diversification remains underexplored. Here, leveraging the adaptive diversity of African Great Lakes’ cichlid species, we combined comparative transcriptomics and population genomics to investigate the evolution of the NC genetic programme in the context of their morphological divergence. Our analysis revealed substantial differences in transcriptional landscapes across somitogenesis, an embryonic period coinciding with NC development and migration. This included dozens of genes with described functions in the vertebrate NC gene regulatory network, several of which showed signatures of positive selection. Among candidates showing between-species expression divergence, we focused on teleost-specific paralogs of the NC-specifier *sox10* (*sox10a* and *sox10b*) as prime candidates to influence NC development. These genes, expressed in NC cells, displayed remarkable spatio-temporal variation in cichlids, suggesting their contribution to inter-specific morphological differences. Finally, through CRISPR/Cas9 mutagenesis, we demonstrated the functional divergence between cichlid *sox10* paralogs, with the acquisition of a novel skeletogenic function by *sox10a*. When compared to the teleost models zebrafish and medaka, our findings reveal that *sox10* duplication, although retained in most teleost lineages, had variable functional fates across their phylogeny. Altogether, our study suggests that NC-related processes – particularly those controlled by *sox10*s – might be involved in generating morphological diversification between species and lays the groundwork for further investigations into mechanisms underpinning vertebrate NC diversification.

## Introduction

The remarkable diversity and complexity of craniofacial structures, pigmentation patterns, and social behaviours within vertebrates is a testament to their outstanding capacity to adapt and exploit a wide range of ecological niches. Much of this phenotypic diversity is intimately connected with the emergence of the neural crest (NC) (Donoghue et al., 2008; Gans and Northcutt, 1983). This embryonic multipotent cell population arises from the dorsal portions of the neural tube and then migrates extensively to finally differentiate into a remarkable range of cell types and tissues, including neurons and glia, pigment cells, craniofacial cartilage and bone, among others (Brandon et al., 2023; Bronner and Simões-Costa, 2016). These diverse cell lineages later assemble to form complex pigmentation patterns in fish, amphibians and birds, as well as divergent head structures, such as fish jaws, bird beaks or mammalian horns (Eames and Schneider, 2005; Elkin et al., 2023; Jheon and Schneider, 2009; Nasoori, 2020).

NC has been primarily studied in the context of its origin, development and function, including developmental disorders involving its derivatives (neurocristopathies) (Bolande, 1997). Studies in model organisms have revealed that the gene regulatory networks (GRNs) and developmental processes governing NC specification, migration and differentiation are highly conserved across distantly related species (Bronner and Simões-Costa, 2016; Simões-Costa and Bronner, 2015). This remarkable macroevolutionary conservation of the NC programme raises key questions about its evolvability and its potential contribution to the origins of vertebrate diversity. Surprisingly, the role that NC cells may play in the the evolution of species-specific traits (i.e. at the population or species level) remains largely unexplored (Abzhanov et al., 2004; Brandon et al., 2023; Donoghue et al., 2008; Kratochwil et al., 2015; Powder et al., 2014). This is despite the rapid and extensive diversification of NC-derived structures - a hallmark of adaptive radiations of multiple vertebrate clades. Striking examples include the diversification of cranial shapes of *Anolis* lizards, beak morphologies in Darwin’s finches and, perhaps most spectacularly, the craniofacial skeletons and colour patterns of cichlid fish radiations in the Great African Rift Lakes (Abzhanov et al., 2006; Albertson and Kocher, 2006; Sanger et al., 2012; Santos et al., 2023).

Here, to investigate the molecular evolution of NC-related phenotypic diversity in the spectacular Lake Malawi cichlid fishes radiation, we first examined the extent of variation in NC genetic and developmental programmes between two closely related, yet eco-morphologically divergent cichlid species, namely *Astatotilapia calliptera* ‘Mbaka’ and *Rhamphochromis* sp. ‘chilingali’. Both species belong to the Lake Malawi cichlid radiation and are characterised by distinct craniofacial morphologies, body pigmentation, ecologies(Bronner and Simões-Costa, 2016) and diets (Edgley and Genner, 2019; Salzburger, 2018; Santos et al., 2023; Turner, 2007) **(Fig. 1a-b** and **Supplementary Fig. 1)**. Our previous work identified variation in pigmentation and craniofacial shapes at the earliest stages of their overt appearance at post-hatching stages (Marconi et al., 2023). Considering the direct mode of development in cichlids (i.e. without larval stage and metamorphosis, unlike zebrafish (Marconi et al., 2023; Woltering et al., 2018), the variation in these NC-derived traits likely stems from differences in early embryogenesis (Marconi et al., 2023) (**Supplementary Fig. 1a-d**).

**Figure 1.**
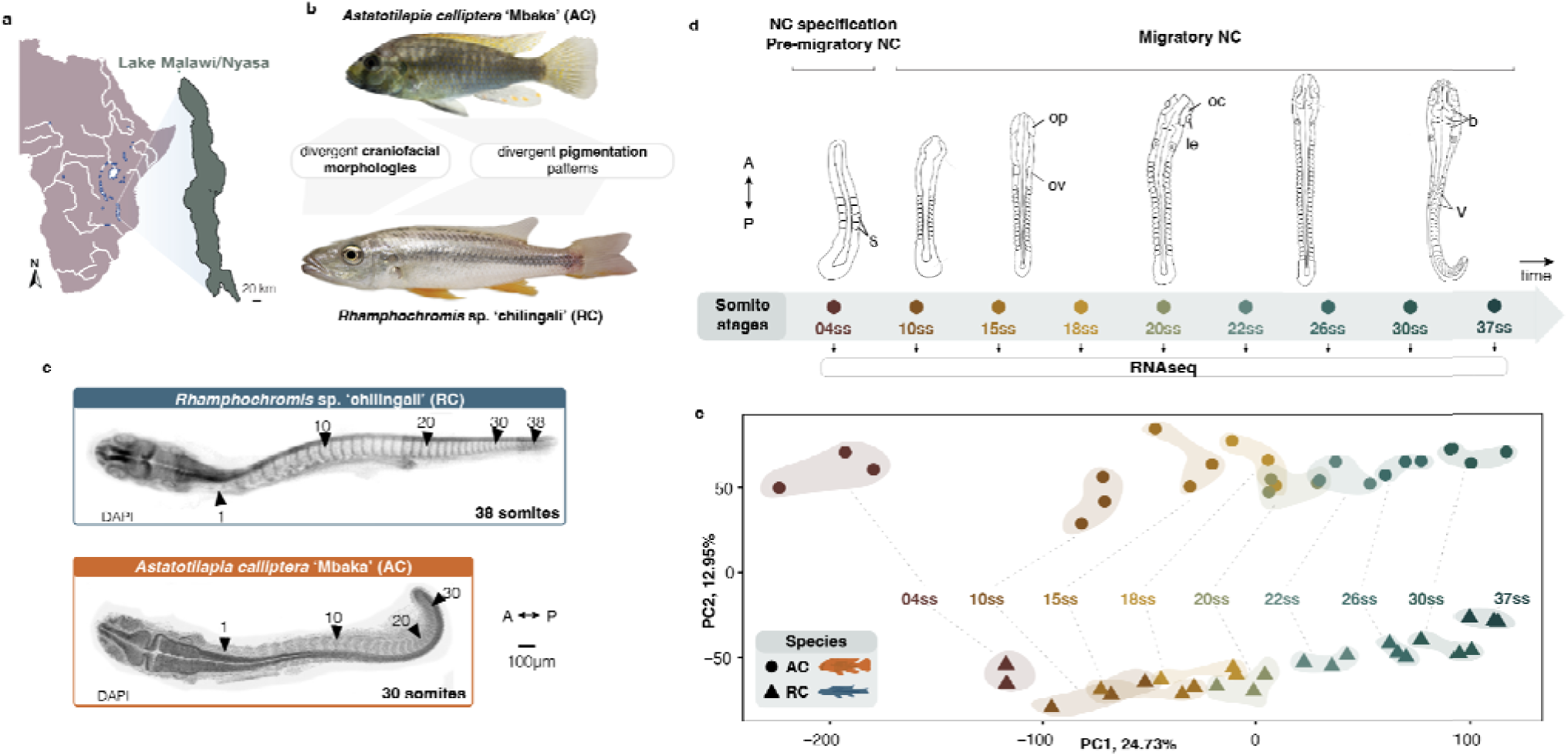
Divergence in whole-embryo transcriptomic trajectories during early embryogenesis and neural crest development between two morphologically distinct Lake Malawi cichlids. **a,** Geographical map of Lake Malawi/Nyasa. **b,** Cichlid species part of this study exhibit distinct neural crest (NC)-derived craniofacial morphologies and pigmentation patterns (to scale). AC has intermediate phenotype of a generalist feeder with variable melanic patches comprising features of both bars and stripes on a yellow-grey background, whereas RC has a flattened head and elongated jaws typical of a piscivore with silvery colouration and dark horizontal stripes (See Supplementary Fig. 1). **c,** AC and RC exhibit different total somite numbers upon completion of somitogenesis (30 in AC, 38 in RC). **d,** The stages of cichlid somitogenesis (expressed as somite stages, ss) examined in this study and collected for RNA sequencing range from the early stages of NC specification (4ss) through migratory NC (10-12ss onwards) and its differentiation during late somitogenesis stages, concluding at 30 and 38ss in AC and RC, respectively (n = 3 biological replicates per ss). **e,** Principal Component Analysis (PCA) of whole transcriptome samples reveals significant ontogenic (PC1) and species-specific (PC2) clustering. Each data point corresponds to a single replicate embryo. A - anterior, AC - *Astatotilapia calliptera* ‘Mbaka’, dpf - days post-fertilisation, le - lens, oc - optic cup, op - optic primordium, ov - otic vesicle, P - posterior, RC - *Rhamphochromis* sp. ‘chilingali’, s - somites, ss - somite stage, st - stage, b - tri- partite brain, V - V-shaped somites. Map (a) modified from d-maps.com.

To test this hypothesis, we focused on the interspecific comparison at earlier embryonic stages concomitant with NC cell specification, migration and onset of their differentiation (Rocha et al., 2020a) (**Fig. 1c-d**). Using whole-transcriptome time-series sequencing data, we uncovered substantial variation in coding and non-coding gene expression throughout NC development between the two species (**Fig. 1d-e** and **Supplementary Fig. 2e-g**) and included divergence in expression levels and temporal trajectories of dozens of genes with key functions within the teleost NC-GRN (Rocha et al., 2020a). Moreover, we show that several of these crucial genes are also associated with signatures of divergent positive selection between species, potentially contributing to species-specific phenotypes. We then focused on two SRY-box transcription factor 10 (*sox10*) paralogs of a key NC specifier, namely *sox10a* and *sox10b*, and showed that they both arose during teleost-specific whole genome duplication and exhibit prevalent inter-specific expression variation throughout NC development. These results suggest potential contribution of *sox10* paralogs, and NC development more broadly, to species differences. Finally, we provide experimental evidence that *sox10a* function is essential for craniofacial skeletal development, indicating a novel role in cichlids that has not been described in any teleost to date. Taken together, our study reveals that *sox10* paralogs followed divergent functional evolution across the teleost phylogeny, including gene loss (zebrafish), subfunctionalisation (medaka) and neofunctionalization (cichlids). We propose that the expansion of genetic toolkit associated with neural crest development during genome duplication, subsequent lineage-specific divergence of paralogous genes, including the acquisition of novel functions, and regulatory and transcriptomic evolution in cichlids, may have collectively contributed to the extensive morphological diversification in this clade. Our results highlight cichlids as a unique teleost system to investigate the developmental and genetic underpinnings of adaptive phenotypic evolution.

## Results

### Genes involved in NC development show species-specific shifts in transcriptomic trajectories

Our previous work has shown that phenotypic divergence between cichlid species in NC-derived craniofacial skeleton and body pigmentation is first observed at the appearance of differentiated cartilage and pigment-bearing cells at early post-hatching stages, respectively (Marconi et al., 2023) (**Supplementary Fig. 1c-d**). Given that cichlids do not undergo metamorphosis and develop directly from embryo to adult (Marconi et al., 2023; Woltering et al., 2018), we hypothesise that variation in processes of NC development occurring early in ontogeny might constitute an important contributor to the adult morphological divergence between these species.

To first investigate the potential role of gene expression variation in driving divergent NC-derived phenotypes, we performed comparative transcriptome profiling (RNAseq) of whole embryos of *Astatotilapia* and *Rhamphochromis* across somitogenesis. This period of embryonic development coincides temporally with NC development (Rocha et al., 2020a) (**Fig. 1c**). In total, 32.49 ± 2.5 Mio paired-end 150bp-long reads were generated for each sample (three biological replicates per somite stage, ss), and then aligned against the *Astatotilapia calliptera* reference genome to quantify gene expression **(Supplementary Table 1** and see **Materials and Methods)**. Principal component analysis (PCA) of protein-coding transcriptomes revealed that gene expression in these species is primarily dictated by ontogeny (i.e. their developmental age in ss; PC1, 24.73%), followed by species (PC2, 12.95%; **Fig. 1e** and **Supplementary Fig. 1e**). Conversely, the expression of non-coding transcripts and transcribed transposable elements (TEs) is primarily clustered by species and then by ontogeny (**Supplementary Fig. 1f-h**), in line with previous reports of higher evolutionary rates associated with non-coding genes in cichlids (El Taher et al., 2021).

We then performed differential gene expression analysis across somitogenesis stages to identify gene candidates showing species- and time-specific transcriptional patterns. In total, 12,611 differentially expressed genes (DEGs; p<0.05) with >1.5-fold expression difference in at least one pairwise comparison were identified (**Fig. 2a, Supplementary Tables 2-3** and **Materials and Methods**). Of these, 14.7% (n=1,857) lacked assigned names in the current assembly *A. calliptera* genome (Ensembl 108), likely representing genes without zebrafish orthologs or novel genes and were excluded from downstream analyses. The remaining DEGs were then classified into seven distinct clusters based on unbiased grouping according to their expression patterns **(Fig. 2a** and **Supplementary Fig. 1i)**. While four of these clusters displayed consistent high or low gene expression in one species across all somite stages (clusters 2, 3, 5 and 6; accounting for 51.4% of all DEGs), the other clusters (1, 4 and 7) showed species-specific temporal shifts in gene expression, possibly linked to the temporal differences (heterochronies) during somitogenesis between these species (Marconi et al., 2023). Each gene expression cluster was significantly enriched for specific Gene Ontology (GO) categories associated with functions ranging from transcription regulation to metabolic and developmental processes (**Fig. 2b**). Notably, clusters 4 and 7, displaying temporal shifts in gene expression dynamics between the two species during early and late somitogenesis, were significantly enriched for genes with functions related to neural crest cell differentiation (**Fig. 2 a-b** and **Supplementary Fig. 1i**). Altogether, the considerable divergence in expression dynamics between species over developmental time (e.g. temporal shifts) may imply variation in multiple developmental processes during embryogenesis, including those involving NC cells. Combined with the diverse repertoire of NC derivatives, these results implicate the potential role of this cell population in the divergence of species-specific traits.

**Figure 2.**
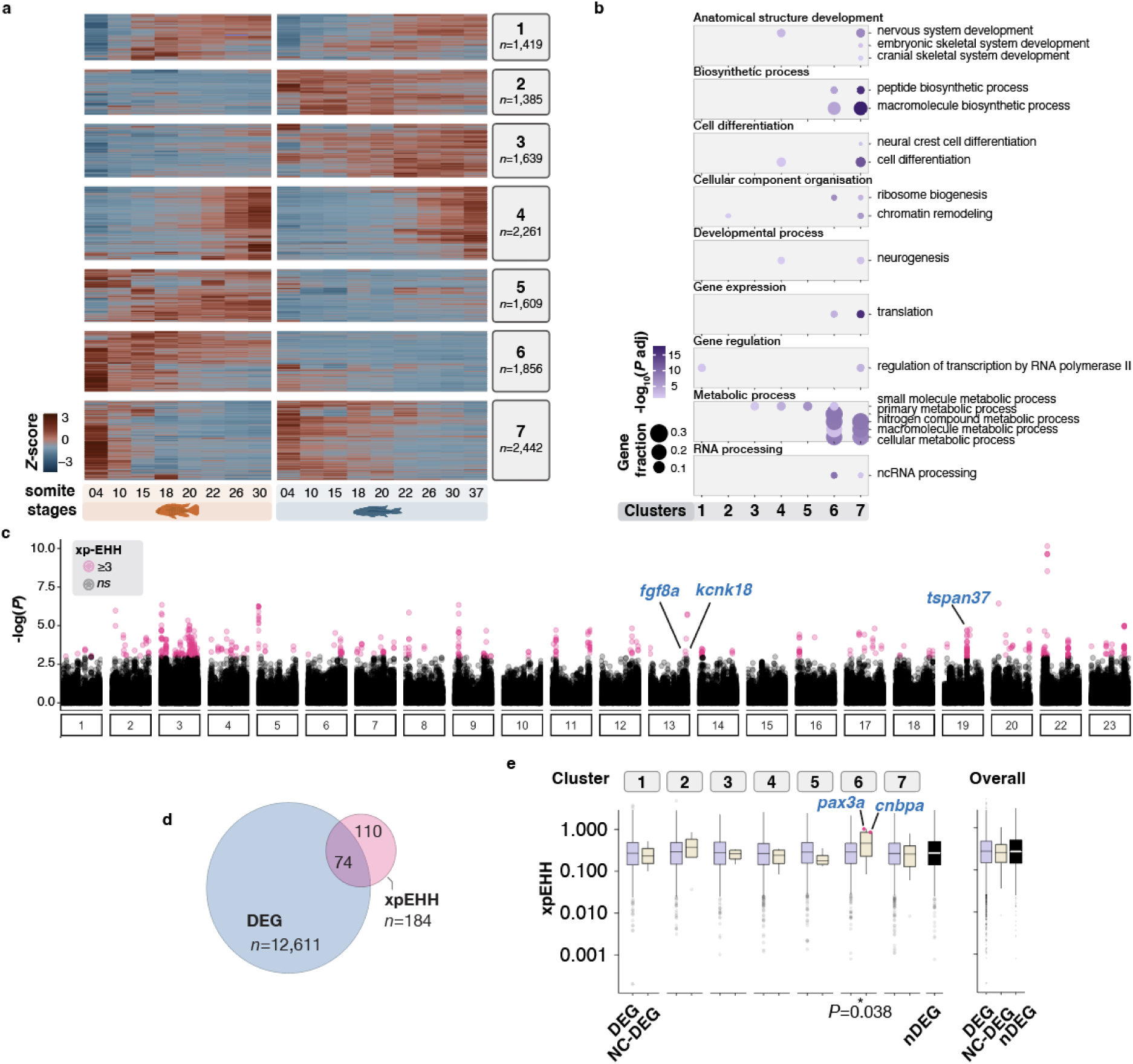
Comparative characterisation of cichlid transcriptomes during somitogenesis and neural crest development. **a,** Heatmap of all differentially expressed genes (DEGs, n=12,611) between any pairwise somite stage comparisons between species identifies seven clusters of gene expression patterns**. b**, Distinct Gene Ontology categories are significantly enriched in each of the seven clusters of gene expression identified. **c**, Genome-wide scans between *Astatotilapia* and *Rhamphochromis* populations reveal 551 significant SNPs (in pink) showing elevated local haplotype homozygosity (measured using between-population extended haplotype homozygosity, xpEHH; see also Supplementary Fig. 2b). **d,** Significant SNPs in close proximity (see Materials and Methods) were grouped into 154 islands, which were associated with 74 DEGs. **e**, NC-DEGs within cluster 6 show significantly elevated xpEHH values, in line with signatures of positive selection. DEG, differentially expressed genes; nDEG, genes lacking significant expression differences; NC-DEG, DE genes linked to NC development.

### Signatures of positive selection are associated with NC-related genes

We then sought to assess if DEGs, and in particular NC-related genes, were potentially diverging between species by conducting genome-wide scans for regions under positive selection. Through extended haplotype homozygosity (xp-EHH, Gautier and Vitalis, 2012) scans between wild *Astatotilapia* and *Rhamphochromis* populations (43-45 whole genomes per species), we identified 154 regions showing significant signatures of positive selection (**Fig. 2c; Supplementary Fig. 2a**, **Supplementary Table 4, Materials** and **Methods**). Altogether, 74 DEGs were located near or within these putative islands of selection **(Fig. 2d**, **Supplementary Fig. 2b-c;** see **Materials and Methods**) with functions ranging from cell differentiation, signal transduction to metabolic pathways, and included several unannotated novel genes (**Supplementary Fig. 2e**).

Among the annotated DEGs located in the most extreme outlier regions were *fgf8a* (cluster 7), *kcnk18* (cluster 1) and *tspan37* (cluster 5), which exhibited significant expression differences between the two species during the early and mid-phases of somitogenesis (**Supplementary Fig. 2b-c**). *tspan37* is an integral membrane protein involved in cellular signalling, while *kcnk18* encodes a potassium channel expressed in the brain and eye of hatchling and larval zebrafish (Rahm et al., 2014). To test whether these genes were involved in NC development and characterise their expression patterns, we performed *in situ* Hybridisation Chain Reaction (HCR) (Choi et al., 2018). Expression of *kcnk18* and *tspan37* was not detected in whole mount embryos at the stages of differential expression, perhaps due to overall low expression levels **(Supplementary Fig. 2c).** Furthermore, *fgf8a*, albeit known for its diverse roles during embryogenesis, including in chondrogenesis of the NC-derived cranial skeleton (Gebuijs et al., 2019), showed expression only in the developing brain and posterior-most notochord during somitogenesis in the examined cichlids **(Supplementary Fig. 3)**, consistent with findings in zebrafish. This limited expression pattern suggests an unlikely role in craniofacial development and patterning at this stage. Altogether, the top divergent outlier genes are not likely to play a role in NC-derived divergence between these two cichlid species. However, NC-related DEGs (identified based on their GO annotation, **Supplementary Table 2, Materials and Methods**) belonging to cluster 6 and showing overall lower expression in *Rhamphochromis* throughout somitogenesis **(Supplementary Fig. 1i)** displayed significant enrichment for sites under potential positive selection (**Fig. 2e** and **Supplementary Fig. 2d**). These included the transcription factor *pax3a*, already implicated in cichlid interspecific pigment pattern variation (Albertson et al., 2014), and the cellular nucleic acid-binding *cnbpa,* involved in craniofacial development in fish (Weiner et al., 2007), among others. Collectively, these findings highlight a strong association between sequence and transcriptional differences between the two species, with several DEGs with functions related to NC processes showing enrichment for sites under potential positive selection.

### Variation across the NC gene regulatory network is particularly associated with NCC migration

Since our comparative transcriptional and selection analyses highlighted divergence in the genetic programme of NC development between species, we next sought to identify genes, processes and stages associated with early NC ontogeny potentially implicated in the evolution of morphological diversity. To this end, we examined only DEGs with known functions (based on their GO annotation, **Supplementary Table 2**) in NC development and differentiation and its two extensively diversified derivatives, namely pigmentation and craniofacial skeleton **(Fig. 3a)**

**Figure 3.**
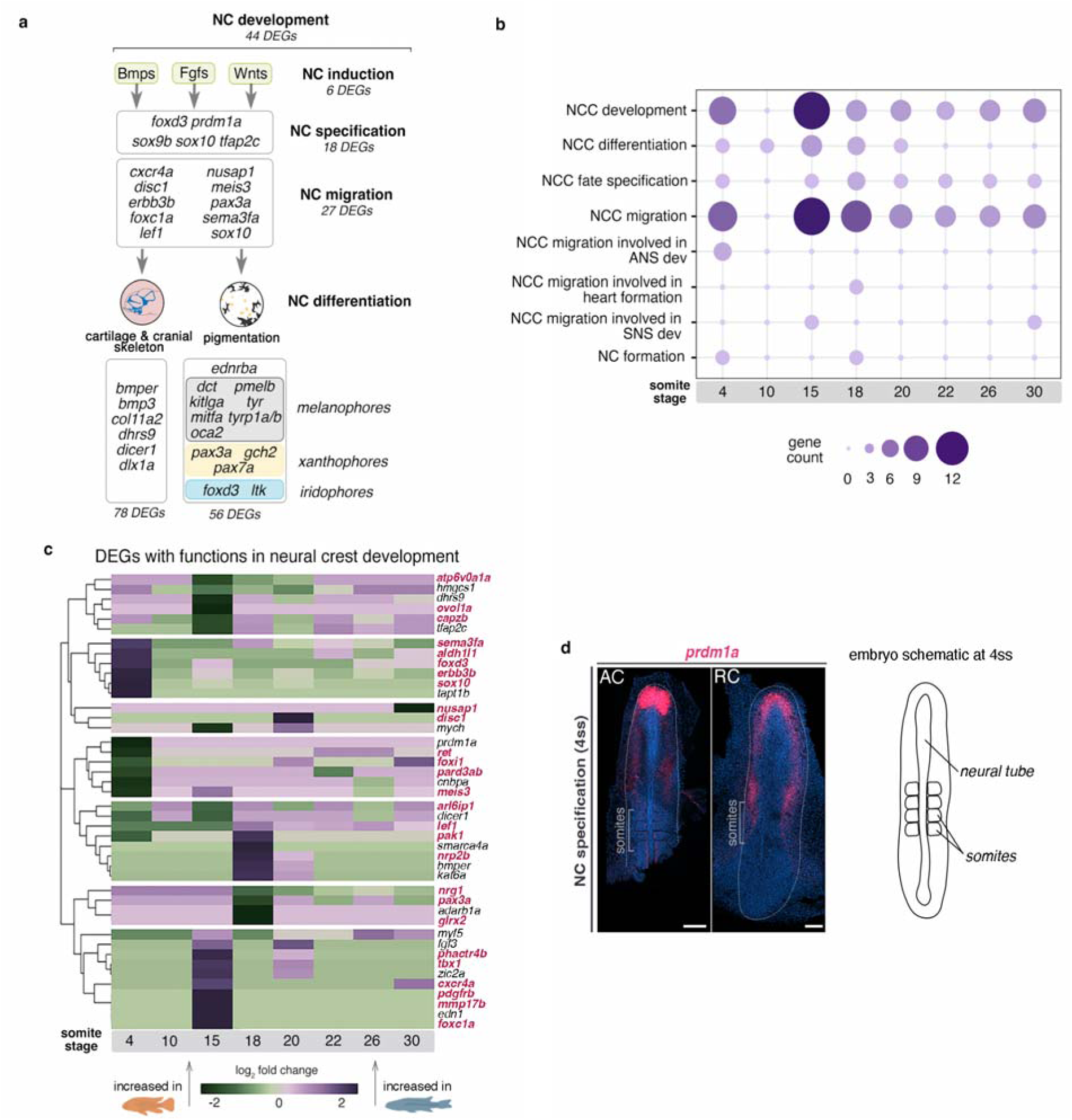
Prevalent transcriptomic variation between cichlids exists across the NC genetic programme. **a,** identified DEGs belong to different tiers of the teleost NC-GRN (Rocha et al., 2020a), from specification to migration and differentiation, including development of NC-derived pigmentation and craniofacial skeleton. **b,** distribution of identified NC-DEGs per function along the RNAseq time-course. c, fold changes in expression levels of candidate genes involved in NC development between cichlids. Genes involved in NCC migration highlighted in red. d, in situ HCR image showing prdm1a expression at 4ss, representative of n ≥ 2 per species. Anterior to the top of the figure. AC - *Astatotilapia calliptera* ‘Mbaka’, ANS - autonomic nervous system, NCC - neural crest cells, RC - *Rhamphochromis* sp. ‘chilingali’, SNS - sympathetic nervous system, ss - somite stage. Scale bar = 100 μm.

The identified NC-DEGs belonged to all tiers of the NC-GRN (Betancur et al., 2010; Sauka-Spengler and Bronner-Fraser, 2008; Simões-Costa and Bronner, 2015) **(Fig. 3a)**. These included genes involved in NC induction, NC specification, NC migration and NC differentiation. This last category included numerous genes associated with the development of different pigment cell lineages, such as melanophores, xanthophores and iridophores (Howard et al., 2021). Multiple genes contributing to the development of the embryonic cranial skeleton, also showed divergence in expression between species as well as four signalling pathways - Bmp, Fgf, Hedgehog and Wnt. Almost 40% of the identified NC-DEGs **(Fig. 3b)** were associated with NCC migration (GO:0001755), and most were differentially expressed at 4ss and 15ss **(Fig. 3b).** These genes (highlighted in red in **Fig. 3c)** exhibited variation in expression levels over time, often displaying large differences in relative expression between species at individual stages, such as 4, 15 and 18ss **(Fig. 3c)**. These findings might reflect divergence in gene expression but also indicate variation in the sizes of cell populations expressing each of these genes (given that the more cells, the more mRNA will be detected in a bulk approach.

The considerable number of DEGs involved in NC migration and differentiation might result from broad knock-on effects of divergence at early stages (i.e. during NC specification) within the NC programme. We identified three candidate genes – *sox10, prdm1a* and *dicer1* – known to perform multiple functions in NC development, including in specification and migration of NC cells and in differentiation of pigment cells and craniofacial cartilages. *sox10* is a key regulator of NC specification, maintenance, migration and differentiation into multiple cell lineages, primarily neuronal and pigment cells, across vertebrates (Carney et al., 2006; Dutton et al., 2001; Kelsh, 2006). *prdm1a* controls NC cell formation by activating *foxd3* (an early NC specifier gene) and regulating *sox10* in zebrafish (Hernandez-Lagunas et al., 2005; Olesnicky et al., 2010; Powell et al., 2013). *dicer1* is required for craniofacial and pigment cell development, and together with miRNAs, it is involved in the regulation of *sox10* during melanophore differentiation in zebrafish (Weiner et al., 2019).

Both *prdm1a* and *dicer1* were differentially expressed at 4ss, a stage concurrent with NC specification (**Supplementary Fig. 2c**), although *dicer1* was not detected in whole-mount specimens. In *Astatotilapia, prdm1a* was highly expressed in the prechordal plate (anterior-most tip of the neural tube) and at lower levels along the neural tube, whereas in *Rhamphochromis*, it was expressed at lower and uniform levels in the prechordal plate on both sides of the anterior neural tube **(Fig. 3d)**. Considering the positive regulatory role of *prdm1a* on *sox10* (Powell et al., 2013), interspecific expression differences in early NC ontogeny could influence later behaviour of migratory NCCs and their differentiation in lineages regulated by *sox10*, such as pigment cells and cartilage. To test this hypothesis, we next investigated in more detail the expression of *sox10* in cichlid embryos to examine the behaviour of migratory *sox10*-labelled NCCs (Drerup et al., 2009; Dutton et al., 2001).

### *sox10* paralogs originated in teleost-specific whole genome duplication while *sox10a* was lost in zebrafish and cavefish

Similar to other members of *soxE* gene family, *sox10* gene is present in two copies (*sox10a* and *sox10b*) in the genomes of cichlids and the majority of other teleosts (Lang et al., 2006; Nagao et al., 2018), with the notable exception of zebrafish – a widely utilised teleost model in biomedical research, which possesses only a single copy, *sox10b* (Braasch et al., 2007; Voldoire et al., 2017). Using recent genomic data of basal teleosts, eels and tarpons (Parey et al., 2023), we confirmed that the *sox10* paralogs indeed originated during the teleost-specific whole-genome duplication event and were subsequently lost in the zebrafish and cavefish lineages **(Fig. 4a)**. Such expansion of the genetic toolkit could provide opportunity for functional divergence, potentially contributing to the process of teleost diversification.

**Figure 4.**
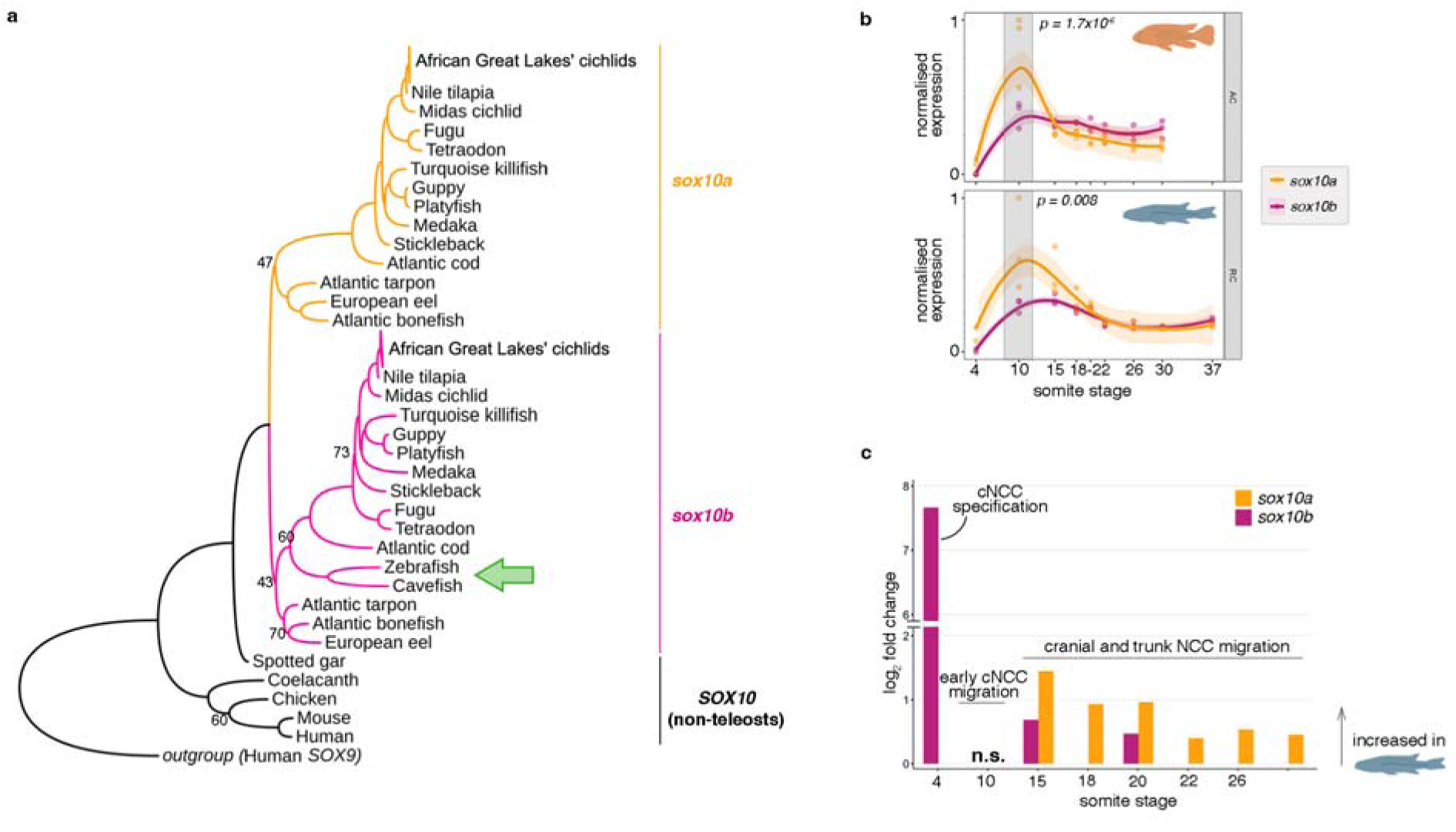
*sox10* paralogs were duplicated in teleosts and are differentially expressed during embryonic development in cichlids. **a,** The topology of Maximum Likelihood phylogeny of *sox10* paralog coding sequences across vertebrates confirms *sox10* duplication occurred at the root of teleosts, followed by *sox10a* loss in zebrafish and cavefish (green arrow). All significant Bootstrap values, apart from the ones shown (<75). **b,** The expression trajectories of *sox10* paralogs across NC development in *Astatotilapia calliptera* ‘Mbaka’ (AC) and *Rhamphochromis* sp. ‘chillingali’ (RC). Shaded bands indicate 95% confidence intervals. Significant differences in normalised gene counts between *sox10* paralogs were observed at 10ss in both examined species (two-way ANOVA and Tukey HSD, *p* < 0.01). **c,** Fold changes in expression levels of *sox10* paralogs between species. Only significant comparisons are shown (*P*-adj < 0.05). cNCC - cranial neural crest cell, NCC - neural crest cell, n.s. - non-significant.

To examine the development of migratory *sox10*-labelled NCCs, we thus focused on both *sox10a* and *sox10b* paralogs. Our transcriptomic profiling revealed that expression of *sox10* duplicates followed similar trajectories over time in both species, with a significant difference in transcript levels between paralogs observed at 10ss (two-way ANOVA and Tukey HSD, p<0.001) **(Fig. 4b)**. Further, the *sox10* paralogs were differentially expressed between species across multiple stages of NC development, with both genes showing consistent upregulation in *Rhamphochromis.* Significant fold expression differences in *sox10b* levels were observed at earlier stages compared to *sox10a* and decreased for both genes with developmental time **(Fig. 4c)**. The differences in expression levels of *sox10* paralogs between and within species during embryonic NC development **(Fig. 4)** suggests that the NC developmental programme, and more specifically NC migration, may be divergent between these two closely related species.

### Differences in expression patterns between *sox10* paralogs suggest divergence in cranial NC development between species

Next, we set out to characterise the precise expression patterns of *sox10a* and *sox10b* in cichlid embryos during the course of NC development. Consistent with findings in medaka (Nagao et al., 2018), the expression of both *sox10* paralogs was generally observed in NC cells at all stages examined, encompassing the processes of NC specification and migration. Furthermore, this expression was detected across different axial levels, corresponding to distinct NC subpopulations (Rocha et al., 2020b), as well as in the otic vesicle (**Fig. 5**). These findings indicate that both cichlid *sox10* paralogs are likely to perform functions in NC development.

**Figure 5.**
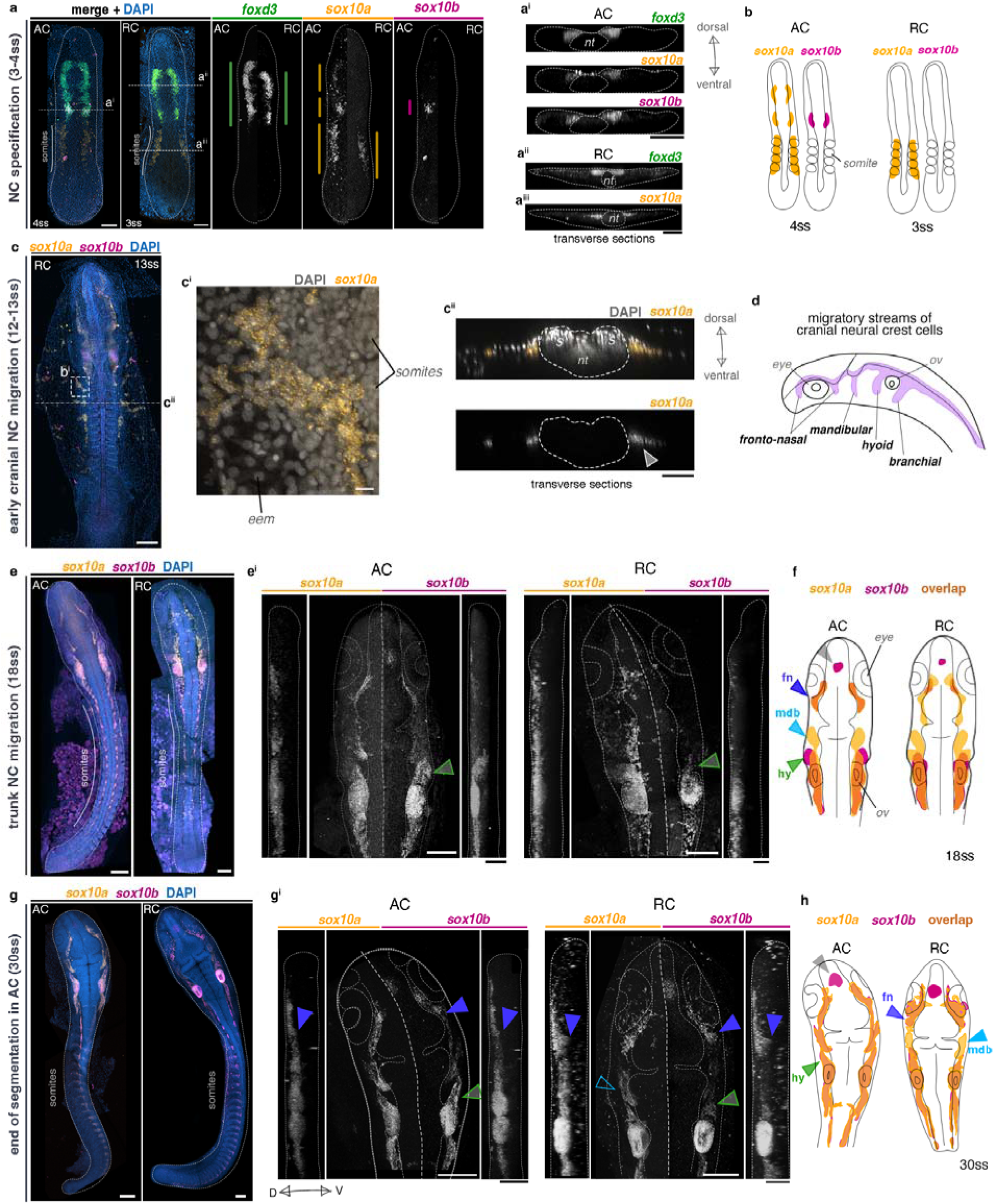
Gene- and species-specific variation in embryonic expression of *sox10* paralogs during neural crest development in Malawi cichlids. *in situ* HCR images of *foxd3* and *sox10* paralogs in representative somite stage-matched embryos of AC and RC. a-a^iii^, early *sox10* paralog expression partially overlaps with *foxd3*, supporting their expression in *bona fide* NCCs, whereas unique somitic domains suggest expression in non-NC cells. B, schematic representation of sox10 paralog expression patterns at NC specification. C-c^ii^, during mid-somitogenesis, *sox10a* is also expressed in extra-embryonic tissues in both species, here shown in RC. d, schematic representation of four migratory streams of cranial NC in cichlids (based on expression of *sox10*) in lateral view. e-h, *sox10* paralog expression in the cichlid embryonic head shows differences between genes and between species, implicating differences in the migratory patterns of NCCs. Schematic representations show expression overlays, highlighting overlapping and unique *sox10a/sox10b*-expressing cell populations. All images present maximum intensity projections of dorsal or lateral (vertical panels in e^i^ and g^i^) views of dissected embryos, with anterior towards the top of the figure. Embryos and sections are outlined in white. Colour-coded vertical bars in a represent the AP extent of expression of each presented gene. Embryos presented in d and e representative of n ≥ 3. AC - *Astatotilapia calliptera* ‘Mbaka’, D - dorsal, NC - neural crest, nt - neural tube, RC - *Rhamphochromis* sp. ‘chilingali’, ss - somite stage, V - ventral. Scale bar = 50 μm, 10 μm in c^i^.

**Figure 6.**
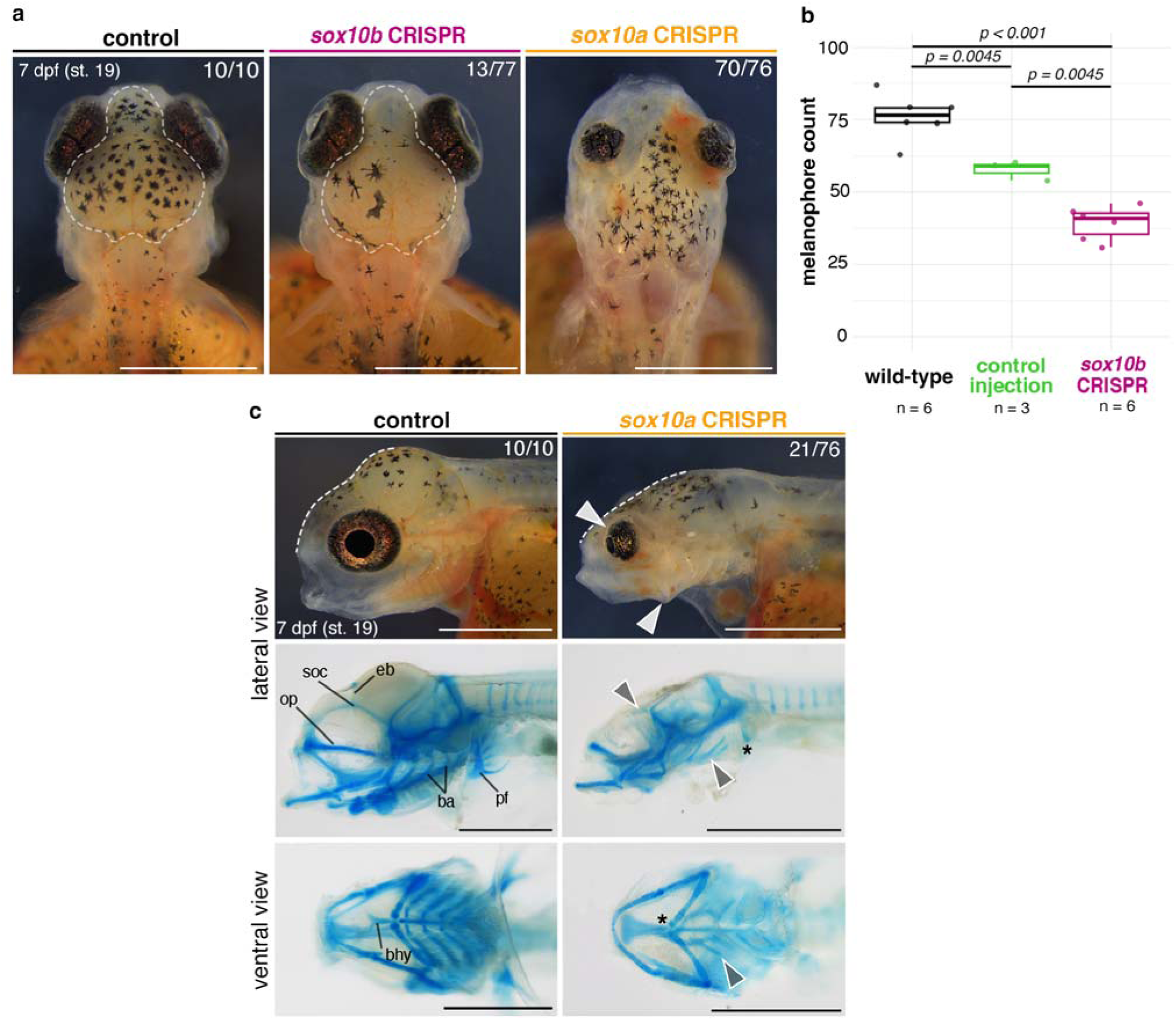
Paralog-specific knockout phenotypes indicate functional divergence of *sox10a* and *sox10b* in cichlids. **a**, Melanophore pigmentation defects are observed in *sox10b* but not in *sox10a A. calliptera* mutants. **b,** *sox10b*-CRISPR embryos have significantly reduced melanophore counts at 7 days post-fertilisation compared to WT and control injections (boxplots showing melanophore count within the dorsal head area outlined in a.; *P*-values are shown for Tukey HSD). **c,** Representative craniofacial phenotypes of *sox10a-*CRISPR embryos in live animals and their corresponding cartilage preparations in lateral and ventral views. White dashed lines show the reduced frontal slope of the brain case. Arrowheads and asterisks in panels showing cartilage preparations indicate reduced and missing cartilages, respectively, in the mutants. Fractions indicate the frequency of presented phenotype across all surviving embryos on that day. Data shown are representative of at least three biological replicates per target gene. Ba - branchial arches, bhy - basihyal, dpf – days post-fertilisation, op - orbital process, soc - super-orbital cartilages, pf - pectoral fin, WT - wild-type. Scale bars = 1 mm.

We identified variation in expression between genes within species during cranial NC specification (4ss) which manifested in temporal and spatial aspects common to both examined species. First, *sox10a* was expressed earlier than *sox10b* and concomitant with *foxd3* (4ss in **Fig. 5a, Supplementary Fig. 4a-b**). Second, *sox10a* was detected in cells residing bilaterally in the dorsal region of somites and that did not co-express neither *sox10b* nor *foxd3* **(Fig. 5a-a^iii^)**. Notably, expression of *sox10a* in the somitic region was also observed at later stages in both species and localised into the extraembryonic membranes extending laterally from the embryo proper enveloping the yolk **(Fig. 5c)**. We propose that these *sox10a+* cells may contribute to the solitary pigmented melanophores that populate yolk during somitogenesis, as they first appear on both sides of the somites in the anterior trunk region at mid-segmentation stages, prior to extensive migration. However, the embryonic origin and function of this novel population remains to be elucidated. Major differences were also observed between species, including multiple *sox10a* domains distributed along the anterior-posterior embryo axis at 4ss in *Astatotilapia* but not *Rhamphochromis* **(Fig. 5a)**. These multiple distinct expression domains of *sox10a*, including in the somitic region, and distinct from those of its paralog *sox10b*, have not been reported for any of the *soxE* family genes in other teleost species to date (Nagao et al., 2018; Takamiya et al., 2020; Tsunogai et al., 2021). Our results thus suggest that *sox10a* could have been co-opted to function in both NC and non-NC cells during early somitogenesis in cichlids, whereas *sox10b* expression during NC specification resembles that of other vertebrates (Aoki et al., 2003; Cheng et al., 2000; Southard-Smith et al., 1998).

As the development progressed, the variation in expression levels and spatial arrangement of *sox10* paralog domains visibly decreased, leading to largely overlapping patterns during cranial and trunk NC migration in both species (**Fig. 5e-h**). We primarily focused on cranial migration, as differences in expression patterns between paralogs and between species were most pronounced in that aspect. During cranial migration, NCCs migrate along highly conserved pathways across all vertebrates, forming four main streams: the anterior-most fronto-nasal stream, followed by the mandibular, hyoid and post-otic branchial stream (**Fig. 5d**) (Steventon et al., 2014). Among these, subsets of cells continued to express only one of the paralogs, for example in *Rhamphochromis* at 30ss, the cells migrating ventrally in the mandibular stream **(mdb on Fig. 5g-h)** expressed only *sox10a* (empty blue arrowhead, **Fig. 5h)**, whereas in *Astatotilapia*, cells migrating in the same stream expressed both *sox10a* and *sox10b* **(blue arrowhead, Figure 5h)**. Furthermore, although both paralogs were expressed in the otic vesicles (but not necessarily co-expressed by the same cells, see **Supplementary Fig. 4c**), only *sox10b* was expressed in a group of cells on the dorsal surface of the forebrain, corresponding to oligodendrocytes derived from neural stem cells (Schebesta and Serluca, 2009) **(grey arrowhead, Fig. 5f and h).** Given the conservation of this pattern across vertebrates (Suzuki et al., 2017), these findings suggest that, compared to *sox10a,* cichlid *sox10b* likely performs a broader range of the conserved vertebrate *sox10* roles in fate regulation of some non-NC derived lineages, such as those derived from neural stem cells.

Several differences were also observed between species in the spatial arrangement and migratory behaviour of the cranial NC cell subpopulations labelled by one or both of the paralogs until the end of somitogenesis in *Astatotilapia* (30ss) **(Fig. 5).** The most pronounced divergence involved the extents of their migration into and around head structures (e.g. otic vesicles, eyes, **Fig. 5)**. For instance, at 18ss, *sox10b+* cells migrating in the hyoid stream **(hy on Fig. 5f and h)**, were present further ventral-laterally in *Astatotilapia* compared to *Rhamphochromis* (green arrows on **Fig. 5e^i^**). This pattern persisted at 30ss **(Fig. 5g^i^)**. In contrast, at the same stage, cells co-expressing *sox10a/sox10b* and migrating in the fronto-nasal stream **(fn on Fig. 5h)** were found further ventrally in *Rhamphochromis* **(purple arrowheads on Fig. 5g^ii^ and h)**.

In summary, cichlid *sox10a* and *sox10b* were expressed by overlapping and divergent subsets of cranial neural crest cells migrating along the stereotypical pathways. Notably, we observed fine-scale variation in migratory patterns of all streams, except for branchial. Such spatial differences could potentially lead to divergence in fine-scale patterning of the structures derived from cranial NCCs i.e. craniofacial cartilages and bones, connective tissues and pigment cells (Kague et al., 2012; Schilling and Kimmel, 1994; Wada et al., 2005). Combined with differences in expression levels from whole-embryo RNAseq **(Fig. 4a-b)**, these differences could be also related to size variation of the populations expressing each paralog, especially considering the differences in embryo sizes between these species (Marconi et al., 2023). Taken together, the variation in expression patterns of *sox10* paralogs in both overlapping and distinct domains indicates divergence in NC development between cichlid species and could reflect potential divergence in developmental functions between two genes.

### Genome editing reveals functional divergence between *sox10* paralogs, including a novel craniofacial skeletal function of *sox10a* in cichlid fishes

Given the extent of divergence between *sox10a* and *sox10b* expression, we set out to characterise their function in cichlids. *sox10* paralog function remains uncharacterised beyond the zebrafish (*Danio rerio*) and medaka (*Oryzias latipes*) model systems. In zebrafish, *sox10b* function is limited to pigmentation and neural derivatives (Kelsh, 2006), while in medaka both paralogs have redundant functions in pigmentation development (Nagao et al., 2018). To test whether *sox10* genes perform divergent roles in cichlids, we deployed CRISPR/Cas9 system in *Astatotilapia* to induce indel mutations in the coding sequence (exon 1) of *sox10a* and *sox10b* in turn **(Fig. 5, Supplementary Fig. 5**-6).

From day 6-7 post-injection (st. 18-19), coinciding with the main stages of craniofacial cartilage development and patterning in this species (Marconi et al., 2023), craniofacial malformations were observed in *sox10a* CRISPR mosaic embryos. Neurocranial and craniofacial deformities were prevalent among injected embryos (**Fig. 5a**, n = 21/76 across four clutches, **Supplementary Table 5**) and, while ranging in severity between clutch-mates, these mutants consistently exhibited flattening of the frontal bones (brain case) (indicated by white dashed lines in **Fig. 5a**), small and bulging, forward-facing eyes as well as protruding, unmoving jaws (white arrowheads in **Fig. 5a**). Alcian Blue stains for cartilage further revealed severely malformed or entirely missing super-orbital cartilages, basihyal, branchial arches and pectoral fins (grey arrowheads and asterisks in **Fig. 5a**). Besides craniofacial abnormalities, *sox10a* mutants at this stage also displayed cardiac and circulatory system defects, reduced black melanophore pigmentation and malformed caudal fin cartilages **(Supplementary Fig. 5)**. The defects in pigmentation were not quantified due to the wide range of severity of cranial deformations, which could have had indirect effects on the pigmentation, for instance due to reduced epidermis surface area for populating chromatophores. The flattened frontal skull slope suggests that the embryonic brains were also likely adversely affected. Mosaic embryos with severe *sox10a*-KO phenotypes **(Fig. 5c** and **Supplementary Fig. 5)** did not survive past 9 dpf (st. 22), suggesting embryonic lethality of a complete KO.

Unlike *sox10*a CRISPR mosaic embryos, *sox10b* mutants did not show any craniofacial cartilage defects at day 7 post-injection (st. 19) nor later, but instead had significantly reduced melanophore pigmentation on the dorsal head region (**Fig. 5b-c**, n = 13/77 across three clutches, **Supplementary Table 5**), the first body area consistently populated by all three differentiated pigment cell types (Marconi et al., 2023). The development of other pigment cell lineages (i.e. reflective iridophores and yellow xanthophores) appeared unaffected by the induced mutations, with no noticeable differences in development of the flank pigmentation at 12 dpf (st. 24) **(Supplementary Fig. 6)**. Despite observed *sox10b* expression in otic vesicles and oligodendrocytes **(Figure 5e-g)** and a known role of zebrafish *sox10b* in glial development (Carney et al., 2006), we did not identify any discernible phenotypes in KO fish involving these tissues.

These functional analyses provide compelling evidence for divergent roles of cichlid *sox10a* and *sox10b* in the development of the neural crest and its derivatives, cartilage and pigment cells (melanophores), respectively. Although we observed a level of potential functional redundancy between paralogs in the differentiation of pigment lineages which requires further investigation, we also uncovered a novel and pivotal role for *sox10a* in the formation of cranial skeleton (neurocranium and craniofacial cartilages) - a function so far only described in cichlids **(Fig. 7)**.

**Figure 7.**
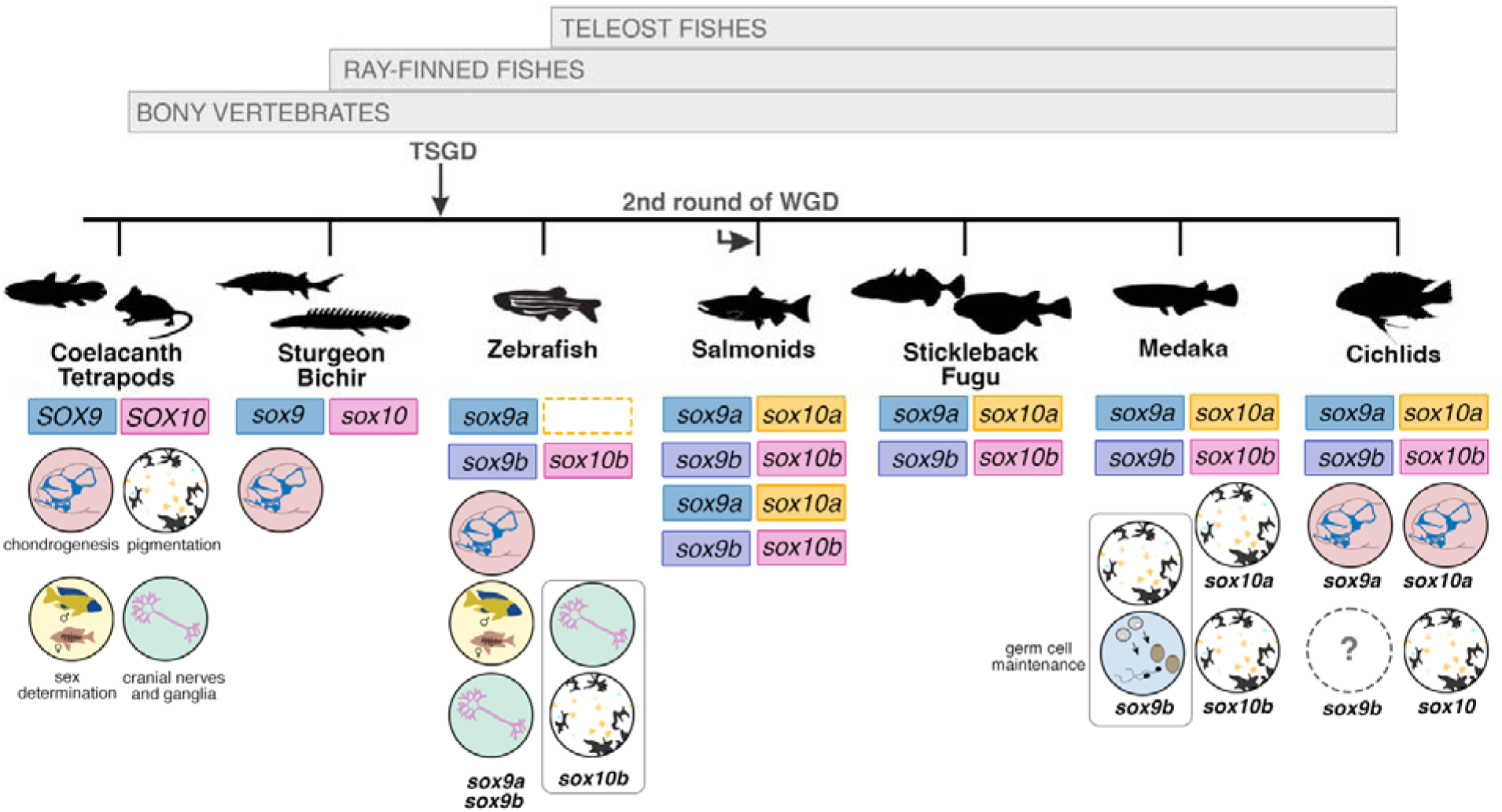
Functional repertoires of *sox10* and *sox9* paralogs across the bony fish phylogeny. Dashed line box indicates gene loss. Bony fish lineage adapted from (Volff, 2005). References for functional analyses are presented in Supplementary Table 6. TSGD - teleost-specific genome duplication; WGD - whole genome duplication. Silhouettes downloaded from http://phylopic.org.

Taken together, the differences in *sox10a* and *sox10b* expression and the *sox10a* role in craniofacial skeletal development show that the cranial NC programme is diverging between cichlid species, but whether the *sox10* paralogs are causally associated with this divergence remains to be tested.

## Discussion

The genetic programme orchestrating development of the neural crest is remarkably conserved across vertebrates, despite NC-derived structures constituting some of the most diverse phenotypic traits, especially among lineages that have undergone adaptive radiation. Our study suggests that neofunctionalization following gene duplication, together with extensive transcriptomic divergence during early NC development, may have collectively contributed to the morphological diversification of NC-derived traits, including pigmentation and craniofacial shapes, in East African cichlids. On a larger evolutionary scale, we report a rare example of taxon-specific, divergent evolutionary trajectories of paralogous genes originating from a single genome duplication event in vertebrates, with *sox10* paralogs showing different fates in different teleost taxa.

### Inter-specific divergence in transcriptional landscape and signature of positive selection in NC-related genes

Recent studies have highlighted that transcriptional evolution among closely-related species and across tissues often underlies phenotypic diversity (Cardoso-Moreira et al., 2019; El Taher et al., 2021). Our findings support and expand upon this concept by uncovering distinct transcriptomic dynamics during somitogenesis between two closely related yet eco-morphologically distinct Malawi cichlid species, with notable differences between coding genes and non-coding and transposable element transcripts. Firstly, we observed that long non-coding genes and TEs displayed predominantly species-specific expression trajectories and presumably higher evolutionary rates compared to protein coding genes. These findings are particularly intriguing and warrant further work, as non-coding genes and TEs have the potential to contribute to phenotypic evolution through multiple mechanisms, including generation of genetic variation, modification of gene regulation and the alteration of genomic architecture (Almeida et al., 2022; Wells and Feschotte, 2020). Although recent case studies have highlighted the roles of TEs in cichlid diversification (Brawand et al., 2014), understanding of their activity during embryonic development remains very limited.

In addition to the non-coding transcriptome, many protein-coding genes, including those involved in the NC development and its derivatives, also exhibited significant variation in expression trajectories between the two species. Notably, differences in expression of many genes could be explained by simple temporal shifts in timing of gene expression relative to somite stage, which in turn can be attributed to differences in developmental timing (i.e. heterochrony) between species during somitogenesis (Marconi et al., 2023). These results provide evidence that variation in developmental timing, and consequently altered gene expression dynamics, contribute to species divergence in this clade.

Using combination of genomic and transcriptomic data, we identified 74 DEGs with signatures of divergent positive selection between species, with the most extreme outliers related to cellular functions, such as *fgf8a* (secreted signalling molecule)*, tspan37* (regulator of cellular signalling) and *kcnk18* (potassium channel protein), and involved in development of several organs and systems, such as brain, eye and nervous system (Gebuijs et al., 2019; Rahm et al., 2014). The diversity of their functions during development makes it challenging to identify a specific phenotype under selection as multiple traits could be involved simultaneously. Further work could examine the functions of these genes in cichlid embryonic development and verify their expression in adult tissues to provide some insights into these results.

Furthermore, our study identified dozens of DEGs with known NC functions, ranging from specification to differentiation into pigment and cartilage cell lineages. Several of these genes have previously been associated with NC-trait variation in cichlid fishes (Albertson et al., 2014) and appear to evolve under divergent positive selection, implying that they might be playing a role in the adaptive evolution of cichlids. The overrepresentation of DEGs involved in NCC migration, including many cell intrinsic (e.g. transcription factors) and extrinsic factors (e.g. signalling molecules), further underscores the causal role that variation in NC migration might play in the emergence of novel NC phenotypes and species differences in cichlids, as posited by studies across vertebrates (Fish et al., 2014; Powder et al., 2014; Tucker and Lumsden, 2004).

### Differences in NC migration between cichlid species potentially associated with trait divergence

Our comparative analysis in two eco-morphological divergent Malawi cichlids revealed that *sox10* paralogs – master regulators within the NC programme – show species-specific temporal and spatial variation, especially in migratory cranial NCCs. This variation could be in turn linked to pigmentation, craniofacial diversity and potentially differences in other NC-derived cell lineages, such as cranial sensory ganglia, Schwann cells and cardiomyocytes (Sande-Melón et al., 2019; Schilling and Kimmel, 1994).

Specifically, *sox10a* had consistently higher expression in *Rhamphochromis* in somite-stage matched embryos. *sox10a+* cells in the fronto-nasal stream showed more advanced in ventral migration and earlier expression in the hyoid stream, suggesting that earlier and, potentially, prolonged migration could lead to larger structures, consistent with the prominent jaws of the piscivore compared to moderate phenotype of an omnivore. However, it is important to note that our *sox10a* mutants displayed relatively normal lower jaw cartilages, while the cartilages of branchial arches (originating from NCCs migrating in the branchial stream) and the frontal slope (fronto-nasal stream) were severely affected. These findings align with patterns of *sox10a* expression in both of these streams throughout cranial NC migration.

Further studies involving later stages (e.g. pharyngula until st. 16 when the first cartilages are present, Marconi et al., 2023) and the use of additional molecular markers (e.g. *sox9* genes, *dlx2a,* collagen gene *col2a1a*) are needed to investigate the potential connection between *sox10a-* expressing cells in the embryo, cartilage formation and these affected structures. Future comparative studies on NC development across multiple divergent species, particularly those characterised by different eco-morphotypes, could provide valuable insights into the connection between NC development, migratory behaviours and adult phenotypes, and how observed differences are associated with their adult adaptive diversity. Nonetheless, our findings highlight the variability of the NC development in closely related cichlids, suggesting unexplored variation in the NC gene regulatory network and highlighting *sox10* paralogs as potential candidates involved in trait diversification.

### Evolution of the NC programme following the teleost genome duplication

One consequence of whole genome duplication (WGD) during teleost evolution is the duplication of entire gene networks. Retained genes post-WGD can contribute to functional innovation and evolution through coding sequence changes and rewiring of the regulatory networks controlling gene expression. The latter is particularly relevant for duplicated regulatory genes, which have a higher potential for significant impacts on gene expression and phenotypic effects (Taylor and Raes, 2004). Our phylogenetic analyses confirmed that *sox10* was duplicated during teleost WGD and retained in duplicate in most teleost genomes, except for zebrafish and cavefish. Interestingly, our expression and functional analyses data revealed distinct fates for *sox10* paralogs in cichlids compared to other teleosts. This presents a unique opportunity to study the impact of gene duplication on NC evolution in multiple evolutionary contexts, including comparisons between non-teleost fishes (i.e. pre-WGD), multiple teleost fishes and other vertebrates.

For example, the differences in expression patterns between *sox10a* and *sox10b* in cichlids are more pronounced than those observed in medaka, particularly with respect to the apparent absence of *sox10a* expression in the extraembryonic tissues in the latter species (Nagao et al., 2018; Tsunogai et al., 2021). The acquisition of novel and divergent expression domains in cichlids suggests the presence of new regulatory elements, warranting future studies to investigate the expression and regulation of *sox10* paralogs in this clade and teleosts on a broader scale.

Moreover, we showed that *sox10a* knockout mutant is associated with aberrant craniofacial skeletal development phenotypes in *A. calliptera*, consistent with the emergence of functional divergence between *sox10a and sox10b* in cichlid fishes and other teleosts. The observed cartilage malformations in mosaic *sox10a*-CRISPR cichlids contrast with the effects of complete knockout mutations of both its orthologs *sox10a* and *sox10b* in medaka and *sox10b* in zebrafish (Kelsh and Eisen, 2000; Nagao et al., 2018). Functional analyses in medaka (Nagao et al., 2018) and zebrafish (Kelsh and Eisen, 2000) suggest that in these lineages *sox10* paralog(s) have retained ancestral functions in pigmentation, neuron and glial cell development, consistent with observations in other vertebrate lineages **(Fig. 7, Supplementary Table 5)**. Notably, medaka *sox10a* and *sox10b* have partially redundant functions in development of pigment cells, in agreement with the most common scenario of subfunctionalization of gene duplicates, also reported for *sox9* paralog in zebrafish (Nagao et al., 2018; Yan et al., 2005). In these teleosts, pigmentation was severely reduced, but cartilage development remained unaffected, similar to other vertebrates (Honoré et al., 2003; Kapur, 1999). The chondrogenesis aberration seen in *sox10a* mutants is more reminiscent of the effects of *sox9a* homozygous mutation in Nile tilapia (Li et al., 2023) and zebrafish (Yan et al., 2005), as well as *sox9* in mice (Wagner et al., 1994) **(Fig. 7, Supplementary Table 5)**.

The expression and functional data combined thus suggest a different partitioning of functions among soxE family genes (specifically *sox9* and *sox10*) in cichlids compared to other teleosts. We posit that *sox10a* acquired an essential role in chondrogenesis in the cichlid lineage, a function performed by *sox9* genes in other teleosts (Cresko et al., 2003; Nagao et al., 2018; Yan et al., 2005), possibly overlapping with the role of *sox9a* in cichlids (Li et al., 2023) **(Fig. 7)**. In contrast, cichlid *sox10b* appears to have retained its function in pigmentation development, akin to its orthologs in other teleost lineages. The subtle, yet significant, pigmentation phenotypes of *sox10b*- CRISPR fish could be due to the mosaic nature of the induced mutations. It remains to be determined whether potential differences in head pigmentation in severe *sox10a* mutants reflect some functional overlap between paralogs or indirect effects of the cranial cartilage malformation on pigment cell development. Similarly, the striking eye and brain size reduction in *sox10a* mutants suggests that chondrogenesis of the craniofacial skeleton might also influence ocular and brain development. Together with observed expression differences of *prdm1a (sox10* regulator) at the anterior neural plate border, these findings may also implicate early variation in cranial placode (developmental precursors of sensory systems) development between species. More comprehensive analyses of NC development in *sox10* mutants, along with examination of non-NC-derived lineages expressing *sox10b,* will be required to better understand the functions of these genes during cichlid embryogenesis. Future studies in a wider range of species will be necessary to assess the extent of functional overlap among other *soxE* family members and to delineate the roles of *sox10a* and *sox9* paralogs in cichlid chondrogenesis, cranial placode and ocular development.

## Conclusion

Functional divergence of duplicated genes is widely recognised for its role in the evolution of morphological diversity, including the expansion of the pigmentation pathway in teleosts (Braasch et al., 2009). In contrast, neofunctionalization is a rare occurrence in genetic evolution, with subfunctionalization (partitioning of ancestral functions between paralogs) being the most common scenario, allowing for developmental fine-tuning (Force et al., 1999). Our findings in cichlids, however, reveal that *sox10a* has acquired a novel function in chondrogenesis, which has not been previously reported in any other teleost clade. This shows that these paralogs have followed divergent evolutionary fates throughout teleost evolution.

We further hypothesise that, while cichlid *sox10b* retained its ancestral function, the neofunctionalization of *sox10a* may have contributed to the rewiring of the NC genetic and developmental programmes. These programmes show remarkable differences between cichlid species, thus providing an opportunity and genetic raw material for the remarkable diversification of the NC-derived phenotypes in cichlids. Altogether, *sox10* paralogs and their variable evolutionary fates are an ideal system for studying the evolution of NC gene regulatory networks across multiple evolutionary scales, within cichlid radiations, and among teleosts and vertebrate species.

## Materials and Methods

### Animal husbandry and embryo culture

Breeding stocks of *Astatotilapia calliptera* ‘Mbaka’ and *Rhamphochromis* sp. ‘chilingali’ were maintained under standardised conditions as previously described in (Marconi et al., 2023). Eggs used for RNA extractions and HCR *in situ* hybridisation experiments were collected from mouthbrooding females immediately after fertilisation and then reared individually in 1 mg/L of methylene blue (Sigma Aldrich) in water in 6-well plates (ThermoFisher Scientific) placed on an orbital shaker moving at slow speed at 27°C until needed. All experiments were conducted in compliance with the UK Home Office regulations.

### Whole embryo bulk RNA sequencing

#### Sample acquisition

For each examined species, all samples were collected from the same egg clutch. Sampling covered the entire period of somitogenesis, which coincides with NC development, with samples taken at 3-hour intervals. At each time point, at least four embryos were dissected and placed individually into either 250 μl of pre-chilled Trizol (Ambion) and stored at -80°C until RNA extraction (at least overnight) or into 1 ml of 4% PFA in 1X PBS for overnight fixation at 4°C. Embryos preserved in 4% PFA were later rinsed twice in 1X PBS and stained with 10 nM DAPI in 70% glycerol in 1X PBS overnight at 4°C, protected from light. Following a wash in 1X PBS (10 min/wash, once), the embryos were mounted on microscopy slides (ThermoFisher) with Fluoromount G (Southern Biotech) and imaged with an Olympus FV3000 confocal microscope to confirm the developmental age (somite stage) of each sampled cohort.

#### RNA extraction

All procedures were conducted on ice, unless otherwise specified. Samples stored in Trizol were thawed from -80°C. For each sample, 100 mg of 0.1 mm zirconia/silica beads (Stratech) were added before homogenization using a TissueLyser II (Qiagen) for 120 seconds at 30 Hz. The samples were then topped up to 1 ml with chilled Trizol and allowed to rest for 5 minutes. Next, 200 μL of chloroform (ThermoFisher Scientific) was added and the samples were vigorously shaken for 15 seconds, briefly vortexed and incubated at room temperature for 15 minutes. The samples were then centrifuged at 300 x*g* for 20 minutes at 4°C. The supernatant was carefully transferred to a fresh tube and further processed using the Direct-Zol RNA Microprep Kit (Zymo) according to the manufacturer’s instructions. The quality and quantity of the extracted total RNA were assessed using Qubit (RNA HS assay, Agilent) and Tapestation (Agilent). Total RNA extracted from each embryo was submitted individually for sequencing, with quantities ranging from 135 ng to 1.3 μg per sample. All sequenced samples had eRIN values above 9.3.

#### NGS library preparation

All libraries were prepared, quality-controlled and sequenced by Novogene Corporation (China) using the Illumina NovaSeq 6000 platform to generate paired-end reads of 150 base pairs (bp). On average, 32.49 ± 2.5 Mio paired-end 150bp reads were generated per sample (**Supplementary Table 1**).

#### Adapter trimming and quality filtering

The adapter sequences in reads were removed, and low-quality sequences (Phred<20) were filtered out with TrimGalore (v0.6.6).

#### Mapping of RNAseq reads to reference genome

All RNAseq reads were mapped to the *Astatotilapia calliptera* genome assembly (fAstCal1.2 in Ensembl 105) using STAR v.2.7.1a (Dobin et al., 2013).

#### Gene expression quantification

The number of reads mapped to each gene in the reference genome was counted in STAR using the built-in HTSeq-count option (Anders et al., 2015). Gene counts were normalised using the median of ratios method in DESeq2 (Love et al., 2014) (v1.34.0). A Principal component analysis (PCA) was applied to reduce the dimensionality of the dataset using the R command prcomp (R 4.2.0).

#### Differential gene expression (DE) analysis and gene annotation

DE analysis was performed on a gene count matrix using DESeq2 (Love et al., 2014) (v.1.34.0). Genes with mRNA counts < 10 per sample in each species were filtered out prior to analysis and technical replicates for each stage were collapsed. Heatmaps of scaled gene expression (*Z*-score per gene across all samples using mean DESeq2-normalised gene count per somite stage per species) were generated using pheatmap (v.1.0.12) and clusters of genes (unbiased complete linkage clustering) were identified and then plotted using ggplot2 (v.3.3.6).

Genes were annotated using the reference genome *A. calliptera* (fAst.cal 1.2) with biomaRt package (Durinck et al., 2009) (v.2.54.0). Significantly differentially expressed genes for each pairwise comparison were identified by filtering for log2 Fold Change above the absolute value of 0.585 (i.e. ≥ 1.5-fold difference in expression) and adjusted p-valueL<L0.05.

#### Gene ontology (GO) analysis

GO annotation and functional enrichment analysis were carried out using gProfiler2 (Kolberg et al., 2020). The full extent of the GO annotation, including the most specific GO terms for each gene or gene product, was used rather than the commonly used *GOslim* annotation, which comprises only a subset of the terms belonging to each parent domain, thus reflecting the broader biological categories. To focus the candidate search on genes involved in the NC development, a dataset comprising all DE genes with GO terms broadly associated with the developmental programme of the NC (including GO terms related to its development, migration and differentiation), development of pigmentation as well as craniofacial complex was compiled (**Supplementary Table 2**). These were then used to explore differences between species.

#### Transposable elements and repeat expression quantification

Transposable elements and repeats were predicted using RepeatModeler (v. 2.0.2 with LTRStruct parameter) and were then annotated using the RepeatModeler custom library in the *A. calliptera* genome using RepeatMasker (v.4.1.4; http://repeatmasker.org/). Only transposon elements (TE) were further analysed (simple repeats, tRNA, rRNA, scRNA and satellites were excluded from subsequent analyses). To quantify TE gene expression, TEcount (v.1.0.1) from the TEtranscript package(Jin et al., 2015) was used following STAR mapping with the following parameters -- chimSegmentMin 10 --winAnchorMultimapNmax 200 --outFilterMultimapNmax 100. Finally, a DESeq2-normalised gene count matrix for 1609 different transcribed TEs was generated and PCA (centred and scaled) was produced with R (prcomp).

### Population genomics

#### Detection of sites under positive selection

To identify recent signatures of positive selections, we conducted genome-wide scans to detect regions with unusually high local haplotype homozygosity and measured the extended haplotype homozygosity between the two populations (XP-EHH) using REHH (v.3.2.2)(Gautier and Vitalis, 2012). We utilised available VCF files containing biallelic SNPs from 43 samples of the Rhamphochromis genus (23.6±8.3x sequencing depth, mean±sd) and 45 randomly chosen *A. calliptera* ‘Masoko Benthic’ (16.02±1.63x) (**Supplementary Fig. 2a**; see Methods in https://github.com/tplinderoth/cichlids/tree/master/callset and (Munby et al., 2021)). Following the approach by (Ravinet et al., 2018), we calculated between-population extended haplotype homozygosity (xpEHH) using phased variants with a minor allele frequency (MAF) >0.05. In total, 551 significant individual sites (SNPs) were identified across all chromosomes (log[*P* value] ≥ 3), suggesting potential positive selection. Significant sites found within 50kbp (±50kbp) were grouped into ‘islands’ using bedTools (2.29.2), resulting in 154 regions, which comprised between 1 and 39 significant SNPs (3.6 SNPs on average) and varied in size between 1bp and 69Mb (mean=4.2kb).

Genes in the closest proximity to these islands were identified using closestBed (bedTools). In total, 184 genes were linked to putative islands of selection (≥1 gene per island), of which 73 were DEGs (**Supplementary Table 4**; 48, 5 and 20 islands located in gene bodies, promoter and intergenic regions, respectively). To identify enrichment in sites under potential positive selection within gene expression clusters, permutation tests were performed between the observed distribution of xpEHH p-values across candidate genes (25kbp regions upstream of TSS) for each gene expression cluster and the expected distribution (over chance). Expected values were calculated by randomly selecting xpEHH values (median values) across size-matched genic windows (1000x iterations; **Supplementary Fig. 2d**).

#### Phylogenetic reconstructions

The coding sequences of *soxE* gene family members were retrieved from Ensembl (108) and directly from genome assemblies from Parey at al. (2023) for inclusion in the phylogenetic analyses (**Supplementary Table 7**). Multiple sequence alignments of *soxE* gene family members were constructed using Clustal Omega (Sievers and Higgins, 2018). Next, the alignments were trimmed with TrimAl (Capella-Gutiérrez et al., 2009) and used to infer evolutionary relationships with the maximum likelihood method in IQ-TREE v.1.6.12 (Nguyen et al., 2015). In-built ModelFinder (Kalyaanamoorthy et al., 2017) was used to infer the best-fit substitution model (TN+F+I+G4) based on the Bayesian information criterion (BIC). The branch supports for maximum likelihood analyses were obtained using the ultrafast boot-strap (UBS)(Kalyaanamoorthy et al., 2017) with 1000 replicates. Phylogenetic tree shown in **Fig. 4** was visualised using iTOL v.6 (Letunic and Bork, 2019) and only non-significant bootstrap values are shown (≤75%).

### Genome editing

#### gRNA design and synthesis

Targets for CRISPR/Cas9 editing were selected with the CHOPCHOP software online (http://chopchop.cbu.uib.no/) (Labun et al., 2019) using the *Astatotilapia burtoni* genome (AstBur1.0) as a reference. Sequence similarity searches with BLAST against the *A. calliptera* genome (AstCal1.2, Ensembl 108) were performed to confirm homology and test for off-target effects. Two sgRNAs targeting exon 1 were designed for *sox10b* (5’-CTCGTCGTCGGATTTGACGG-3’ and 5’-CGCGGATTCCCGCGGGGAA-3’) and *sox10a* (5’-CGGTCAGTCAGGTGCTGGACGGG-3’ and 5’-TCGTTTCCCGATCGGCATAA-3’), respectively, and purchased from Integrated DNA Technologies (ITD) as Alt-R CRISPR-Cas9 sgRNA (2 nmol).

#### Microinjection

Single-cell embryos of *Astatotilapia calliptera* ‘Mbaka’ were injected and maintained following the protocol described in (Clark et al., 2022). We were unable to perform microinjections in a similar m in *Rhamphochromis* due to their prolonged and unpredictable breeding behaviour, rendering it technically unfeasible.

#### Embryo imaging

Injected and control embryos were imaged daily until 12 days post-fertilisation using a Leica M205 stereoscope with a DFC7000T camera under reflected light darkfield. All specimens were positioned in 1% low melting point agarose (Promega) and anaesthetised with 0.02% MS-222 (Sigma-Aldrich) if required to immobilise during imaging.

#### Genotyping

Tissue samples were taken from specimens sacrificed by overdose of 0.5% MS-222 (Sigma-Aldrich) to extract genomic DNA using PCRBIO Rapid Extract Lysis Kit (PCRBiosystems). Fragments of 190-420bp surrounding predicted deletion sites were amplified using PCRBIO HS Taq Mix Red (PCRBiosystems) with an annealing temperature of 56°C using following primer pairs: *sox10b* 5’- CTGTCACCGGGTCATTCCTC-3’ and 5’-GCGTTCATTGGCCTCTTCAC-3’; *sox10a* 5’- ATGGTCACTCACTGTCACCG-3’ and 5’-CCTCCTCGATGAATGGCCTC-3’. Amplicons were purified with QIAquick PCR Purification Kit (Qiagen) before Sanger sequencing. All protocols were conducted following manufacturer’s instructions. Sequence analysis to infer CRISPR edit sites was performed using the Synthego ICE CRISPR analysis tool (https://ice.synthego.com/).

### Cartilage preparations

Embryos were stained for cartilage following the protocol of (Marconi et al., 2023) with following modifications: (1) all specimens were bleached to remove melanophore pigmentation using a solution of 0.05% hydrogen peroxide (Sigma) and 0.05% formamide (ThermoFisher) for 30-45 mins under light and (2) samples were cleared using first 50% then 70% glycerol:water solutions until complete sinking. Specimens were stored in 70% glycerol until imaging in 80% glycerol using a Leica M205 stereoscope with a DFC7000T camera under reflected light.

### Whole mount *in situ* hybridisation by Hybridisation Chain Reaction (HCR)

#### Reagents

The HCR probes and hairpin sets were ordered from Molecular Instruments, whereas all required buffers were made following the instructions provided by the manufacturer. The HCR probe sets (14-20 pairs per gene) were designed using target gene template sequences retrieved from *A. calliptera* genome assembly (fAstCal1.2, Ensembl 108) (**Supplementary Table 8**). Each probe set was designed by the manufacturer to target transcript regions common to all splicing isoforms while minimising off-target effects.

Due to a very low genetic variation in the coding sequences between study species, we used the same probe sets per each target gene for both cichlid taxa examined. Probe specificity was verified by BLAST searches against the *A. calliptera* genome available on Ensembl (AstCal 1.2) as well as against unpublished *Rhamphochromis* sp. ‘chilingali’ assembly.

#### Protocol overview

Dissected embryos were fixed overnight at 4°C in 4% PFA in 1X PBS. The following day, they were rinsed twice in 1X PBST (1X PBS+0.01% Tween-20) without incubation and washed in 1X PBST (10 min/wash, twice) before a stepwise dehydration to 100% MetOH in pre-chilled solutions of 25%, 50% and 75% MetOH:PBST (10 mins/wash, at 4°C) and stored at -20°C until further analyses.

The embryos were then rehydrated from 100% MetOH to 1X PBST in reciprocal series (5 mins/wash, at 4°C), followed by washes in 1X PBST at room temperature (5 min/wash, twice).

mRNA *in situ* hybridization by chain reaction (HCR) was carried out according to the protocol of Andrews *et al*. (2020)(Andrews et al., 2021) for whole mount amphioxus embryos with the following modifications. (1) 2 pmol of each probe mixture (1 μL of 2 μM stock) per 100 μL of probe hybridization buffer were used and (2) 60 pmol of each fluorescently labelled hairpin (i.e. 2 μL of 3 μM stock) were applied per 100 μL of amplification buffer. Finally, the embryos were stained with 10nM DAPI in 70% glycerol in 1X PBS overnight at 4°C protected from light, washed in 1X SSCT (10 min/wash, twice) before mounting with Fluoromount G (Southern Biotech) on glass bottom dishes (Cellvis) without coverslip or microscopy slides (ThermoFisher Scientific) with #1.5 coverslips (Corning), depending on the size of the specimen. To prevent embryos from getting squashed when mounting on slides, thin strips of electrical tape were used as bridges to create space between the slide and a coverslip. Clear nail varnish was used to seal the edges of the slide and all samples were cured overnight at room temperature protected from light before imaging.

All *in situ* hybridization experiments were performed with multiple specimens from different clutches (at least 3 individuals per clutch, repeated at least once with specimens from alternative clutches) to fully characterise the expression patterns.

### Confocal microscopy

Imaging of dissected and stained embryos was carried out with an inverted confocal microscope Olympus FV3000 at the Imaging facility of the Department of Zoology, University of Cambridge. Since the fluorescence intensity levels were only compared as relative signals within each sample (i.e., embryo), imaging was performed using optimal laser power and emission wavelength for each sample. Sequential acquisition mode was used to minimise the signal crosstalk across channels and all images were acquired at 1024x768 resolution and 12-bit depth.

### Image processing for figure presentation

Confocal micrographs were stitched using the Olympus FV3000 software and processed with Fiji(Schindelin et al., 2012) to produce optical sections, collapse z-stacks and adjust image brightness and contrast where necessary following guidelines by Schmied & Jambor (2021)(Schmied and Jambor, 2021). The presented transverse optical sections are maximum projection across 5 adjacent slices. Auto-fluorescence and background noise in each image was removed by subtracting the average pixel intensity measured for each channel in regions of the embryo where no fluorescent signal was observed. Any remaining overexposed pixels were removed using the “Remove outliers” (radius = 0.5 pixel, threshold = 50) function in Fiji. Images presented as figures were smoothened by applying a Gaussian filter with σ = 0.5. Image look-up tables (LUTs) were taken from the BIOP plugin (https://imagej.net/plugins/biop-lookup-tables) and “ChrisLUTs” (Christophe Leterrier and Scott Harden; github.com/cleterrier/ChrisLUTs) package for Fiji/ImageJ. The images were processed for any background imperfections and assembled into figures in Adobe Photoshop 2023.

All plots and statistical analyses were made using R version 4.1.1. The MetBrewer package (https://github.com/BlakeRMills/MetBrewer) was used for colour palettes throughout this work.

## ACKNOWLEDGMENTS

AM was supported by Wellcome Trust (WT) PhD Programme in Developmental Mechanisms ((215223/Z/19/Z), EMS is supported by a Natural Environment Research Council (NERC) Independent Research Fellowship (NE/R01504X/1) and GV is supported by an EMBO fellowship (ALTF 339-2022). The authors thank the Department of Zoology Imaging Facility for assistance with microscopy, and Dr. H. Svardal (University of Antwerp), Dr. G. Turner (Bangor University), Dr. D. Joyce and A. Smith (University of Hull) for the kind gifts of fish stocks and samples. We thank Dr. Ben Steventon and the Santos lab for critical comments on the manuscript. For open access, the authors have applied a CC BY public copyright licence to any author-accepted manuscript version arising from this submission.

## AUTHORS’ CONTRIBUTIONS

AM, GV and EMS devised the study and analyses. MJN collected wild specimens to establish laboratory stocks. RD contributed to whole-genome data collection. AM collected and performed RNA extractions on the samples. AM and GV performed data analyses on sequencing data. AM and AK collected embryonic material, performed HCR staining and imaging. AM performed CRISPR/Cas9 mutagenesis, embryo imaging, phenotyping, cartilage staining and DNA collection. AM, GV and EMS wrote and revised the manuscript. All authors read and approved the final manuscript.

## COMPETING INTERESTS

The authors declare no competing interests.

## INCLUSION & ETHICS

The Malawi cichlid samples (*Rhamphochromis* species) were collected ethically under prescribed permits, and the results and data are published under an Access and Benefit Sharing agreement with the Government of Malawi. We acknowledge the contributions of the Malawi Department of Fisheries and the Government of Malawi for their assistance in the collection of samples and the generation of data and results.

## DATA AVAILABILITY

The raw RNA sequencing reads have been deposited in the Gene Expression Omnibus and will be made public upon publication. This data is made available on an open access basis for research use only. Any person who wishes to use this data for any form of commercial purpose must first enter into a commercial licensing and benefit sharing arrangement with the Government of Malawi.

## CODE AVAILABILITY

All scripts used to analyse the data can be accessed at: https://github.com/Santos-cichlids/neural-crest-sox10-paralogs-cichlids

## Notes

https://github.com/Santos-cichlids/neural-crest-sox10-paralogs-cichlids

### Competing Interest Statement

The authors have declared no competing interest.

### Summary of Updates

Main text figures and sections of results and discussion were revised to clarify the findings and conclusions of the paper

