## Supplementary Table 6 for "Genetic and developmental divergence in the neural crest programme between cichlid fish species"

**Supplementary Table 6.** Summary of *sox9* and *sox10* paralog mutant phenotypes reported in published literature.

|  | ***sox9a*** | ***sox9b*** | ***sox10a*** | ***sox10b*** |
| --- | --- | --- | --- | --- |
| Lake Malawi cichlids *Astatotilapia calliptera* and *Rhamphochromis* sp. 'chilingali' (this study) | / | / | Craniofacial cartilage, cardiac, ocular and pigmentation defects | Pigmentation defects |
| Nile tilapia (*Orechromis niloticus*) | Cartilage defects^1^ | / | / | / |
| medaka (*Oryzias latipes*) | / | Pigmentation defects^2^ | Pigmentation defects^3^ | Pigmentation defects^3^ |
| Zebrafish (*Danio rerio*) | Cartilage defects^4^ | Cartilage defects^4^, peripheral neuron and glial defects^5^ | NA | Pigmentation defects6^,^ peripheral neuron and glial defects^7-9^ |
|  | ***SOX9*** | | ***SOX10*** | |
| Mouse (*Mus musculus*) | Skeletal malformations^10^ | | Aberrations of cranial nerve and ganglia morphology^11^ | |

/ - mutant phenotype unknown

NA – gene absent

1 - Li et al., 2023 *Zoological Research*

2 - Tsunogai et al., 2021 *Development, Growth & Differentiation*

3 - Nagao et al., 2018 *PLoS Genetics*

4 - Yan et al., 2005 *Development*

5 - Carney et al., 2006 *Development*

6 - Kelsh 2000 *BioEssays*

7 - Kelsh and Eisen, 2000 *Development*

8 - Elworthy et al., 2000 *Mechanisms of Development*

9 - Delfino-Machin et al., 2017 *PLoS ONE*

10 - Wagner et al., 1994 *Cell*

11 - Kuhlbrodt et al., 1998 *Journal of Neuroscience*
