## Supplementary Fig. for "Genetic and developmental divergence in the neural crest programme between cichlid fish species"

**Marconi, Vernaz et al.**

**SUPPLEMENTARY MATERIALS**

- **Supplementary Tables**
- **Supplementary Table 1.** Information regarding RNAseq.

*Supplementary_Table_1_SeqRead.xlsx*

- **Supplementary Table 2**. Gene Ontology terms related to neural crest, pigmentation and craniofacial skeleton development used to subset DEGs.

*Supplementary_Table_2_candidateGOterms.xlsx*

- **Supplementary Table 3.** Gene count matrix for DEG candidates associated with Gene Ontology of interest (from Supplementary Table 2).

*Supplementary_Table_3_DEG_candidates.xlsx*

- **Supplementary Table 4.** List of DEGs associated with xpEHH outlier peaks.

*Supplementary_Table_4_xpEHH_DEG.txt*

- **Supplementary Table 5.** Sample sizes for CRISPR/Cas9 experiments.

*Supplementary_Table_5_CRISPR_stats.xlsx*

- **Supplementary Table 6.** Summary of mutant phenotypes for *sox9* and *sox10* genes in bony fishes.

*Supplementary_Table_6_Summary_mutant_phenotypes.docx*

- **Supplementary Table 7.** Accession numbers for *sox10 orthologs and sox9* coding sequences used for phylogeny reconstruction.

*Supplementary_Table_7_sox10_sox9_accessions.xlsx*

- **Supplementary Table 8.** Accession numbers for coding sequences used to design HCR probes targeting candidate genes and their lot numbers.

*Supplementary_Table_8_HCRprobes.xlsx*

**Supplementary Figures**

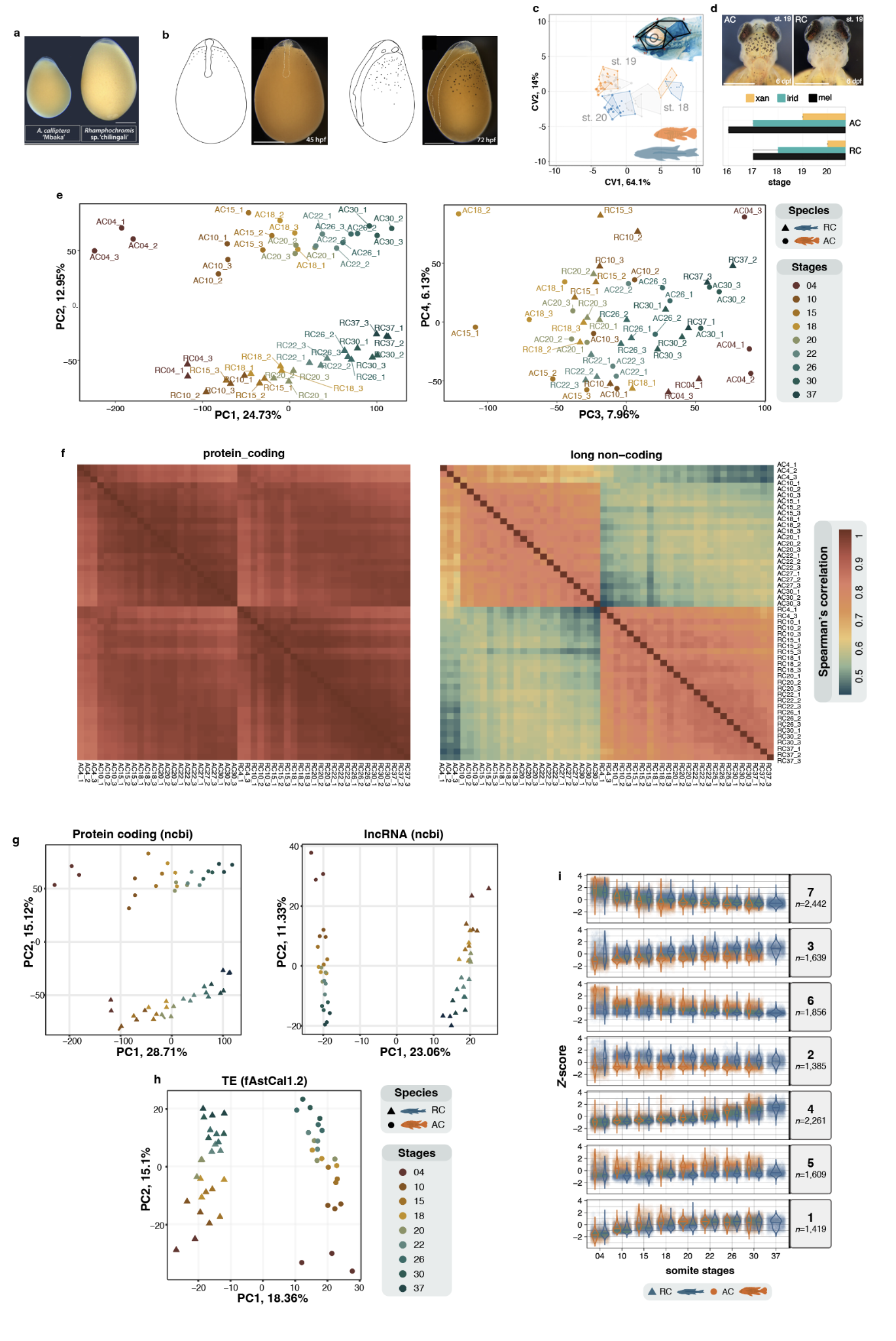

**Supplementary Figure 1. Comparative characterisation of cichlid transcriptomic landscapes during somitogenesis and neural crest development.** **a,** Images of fertilised eggs for AC (left) and RC (right). **b,** Example of embryo dissected from the yolk. **c, d**, Interspecific divergence in shapes of the craniofacial skeleton (depicted in a common morphospace in **c**) and timing of pigment cell appearance (**d**) during post-hatching development (stages 16-20) indicates that phenotypic variation in these NC-derived traits is specified prior to their overt formation (Marconi et al., 2023)**. e**, PCA plots showing PC1-PC2 (left) and PC3-PC4 with labelled samples (right) as part of the whole transcriptome analysis (see Fig.1e). **f.** Heatmaps showing pairwise Spearman’s correlation scores of gene expression values for protein coding (23,664 transcribed genes with≥5 normalised count in any one sample) and non-coding transcripts (1664 transcribed genes with ≥5 normalised count in any one sample). **g-h**, PCA plots of gene expression values (DESeq2 gene count normalisation) for protein coding and lncRNA (**g.**), and for transcribed TE transcripts (1,609 TE transcripts) (**h.**). **i,** Gene expression dynamics (*Z*-score, scaled normalised gene count) for all DEGs across all somite stages according to seven gene expression clusters identified in Fig. 2a. Hpf, hour post-fertilisation. AC, *Astatotilapia calliptera* ‘Mbaka’; RC, *Rhamphochromis* sp. ‘chilingali’.

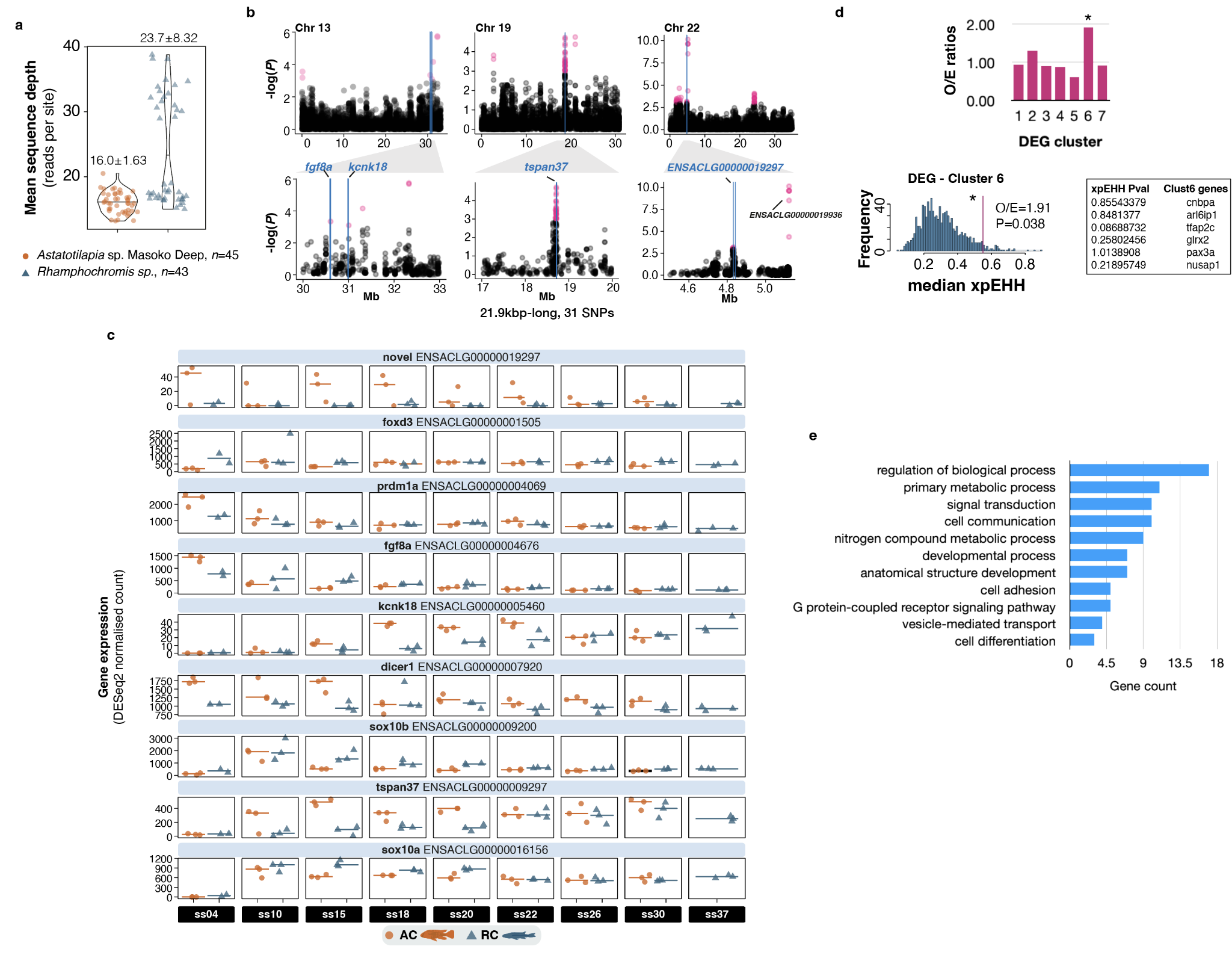

**Supplementary Figure 2. Comparative population genomics between AC and RL cichlid species identifies regions of positive selection associated with transcriptomic divergence. a**, Violin plot showing overall genome coverage for the two cichlid population used genomics analyses (horizontal bars for median values). **b,** Close-up genome-browser views focusing on the DEG candidates associated with xpEHH peaks shown in Fig.2c. **c,** Plots of expression values for some DEG candidates associated with outlier regions of putative signatures of selection (refer to Fig. 2c); horizontal bars show median values. **d.** Upper panel: observed vs. expected ratios of xpEHH significance enrichment for all the NC-DE genes belonging to each DEG cluster. Expected values were computed through 1000 random iterations. NC-DE genes from cluster 6 show significant enrichment for xpEHH significance, indicative of signature of positive selection. Lower panel: O/E distributions of xpEHH for NC-DEG belonging to cluster 6 (listed on the left). **e.** Top 10 GO categories for DEGs associated with xpEHH outlier peaks.

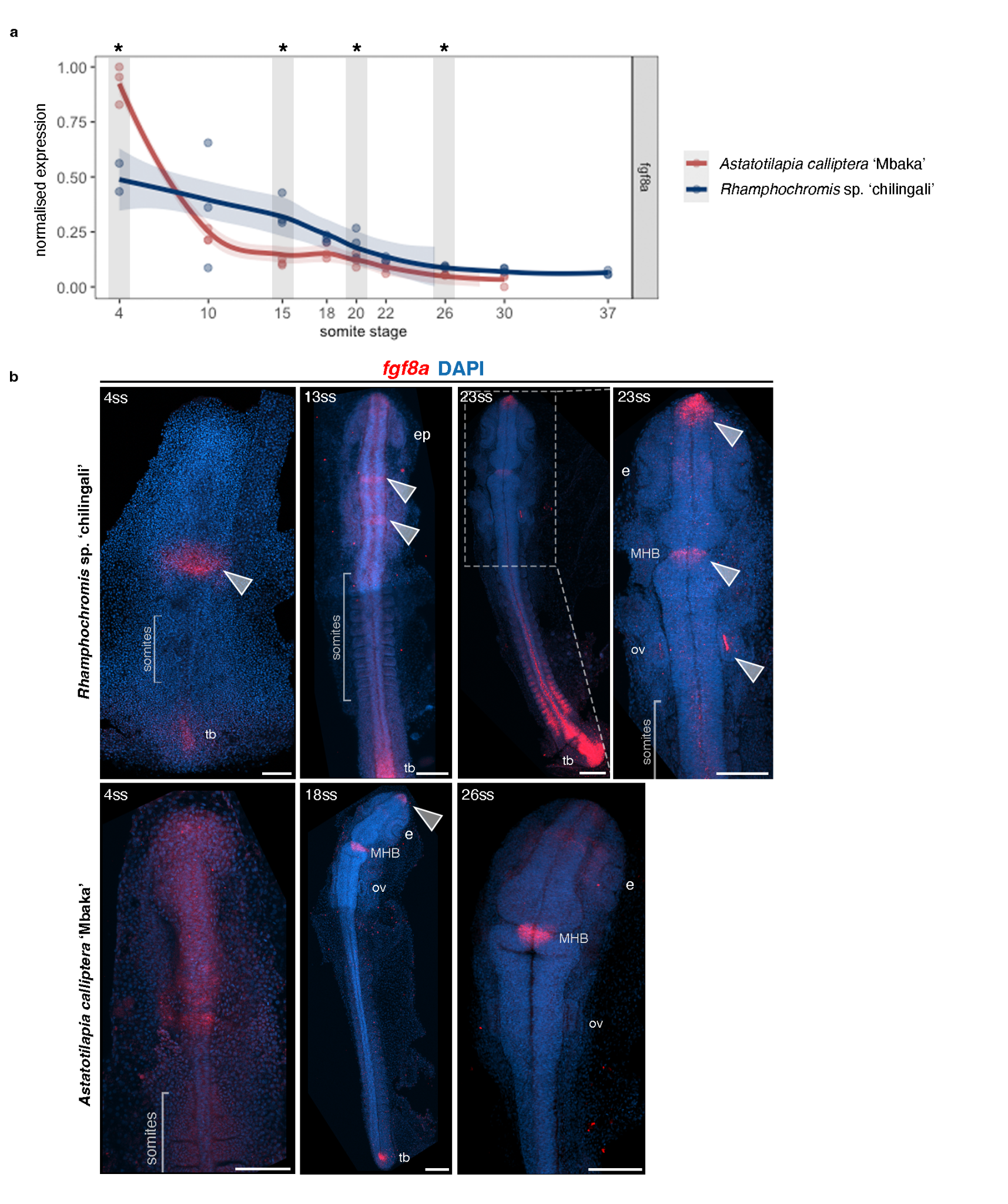

**Supplementary Figure 3. *Fgf8a* expression patterns in cichlid embryos suggest a role in brain development during somitogenesis rather than neural crest.** **a**, Significant differences in expression levels of *fgf8a* were observed at multiple stages of somitogenesis. Shaded panels and asterisks denote stages of differential expression between species with log_2_ fold change over 1.5 and p-adj < 0.05. **b**, Expression patterns of *fgf8a* in cichlids at selected stages of somitogenesis. Note the difference in expression domains between species at 4ss. At later stages, in addition to strong expression at the midbrain-hindbrain boundary (MHB) and anterior-most end tip of the forebrain, *fgf8a* transcripts were also detected in the tailbud (arrowheads). Expression in otic vesicles was only observed in *Rhamphochromis*. ep - eye primordium, e - eye, MHB - midbrain-hindbrain boundary, ov - otic vesicle, tb - tailbud. Scale bar = 100 μm.

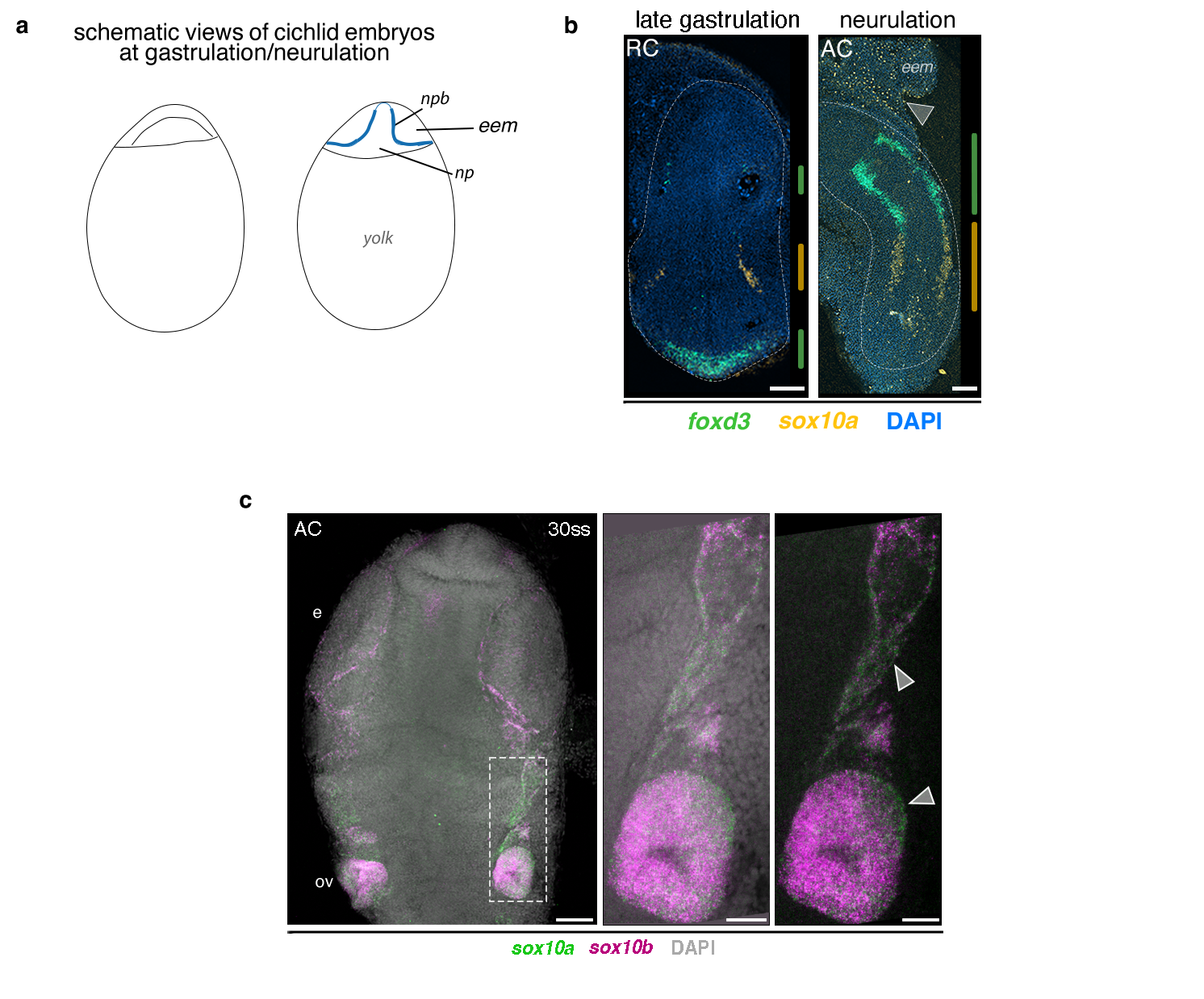

**Supplementary Figure 4. Temporal divergence between *sox10* paralogs and novel expression domain of *sox10a*. a-b,** In embryos undergoing gastrulation (a and b left) and neurulation (a and b right), *sox10a* expression was detected prior to *sox10* in domains distinct from *foxd3*, a marker of early neural crest cells. **c,** *sox10a* and *sox10b*, although largely co-expressed in neural crest cells during somitogenesis, were also detected in distinct nuclei in cranial neural crest, indicating paralog-specific cell populations (grey arrowheads). *sox10a* depicted in green to improve visual distinction of HCR signals. Note that, unlike zebrafish, epiboly in cichlids is not complete until late segmentation/early pharyngula stages (Marconi et al., 2023). AC - *Astatotilapia calliptera* ‘Mbaka’, eem - extra-embryonic membrane, np - neural plate, npb - neural plate border, RC - *Rhamphochromis* sp. ‘chilingali’, ss - somite stage. Scale bar = 100 μm, 50 μm in c left panel, and 25 μm in c central and left panel.

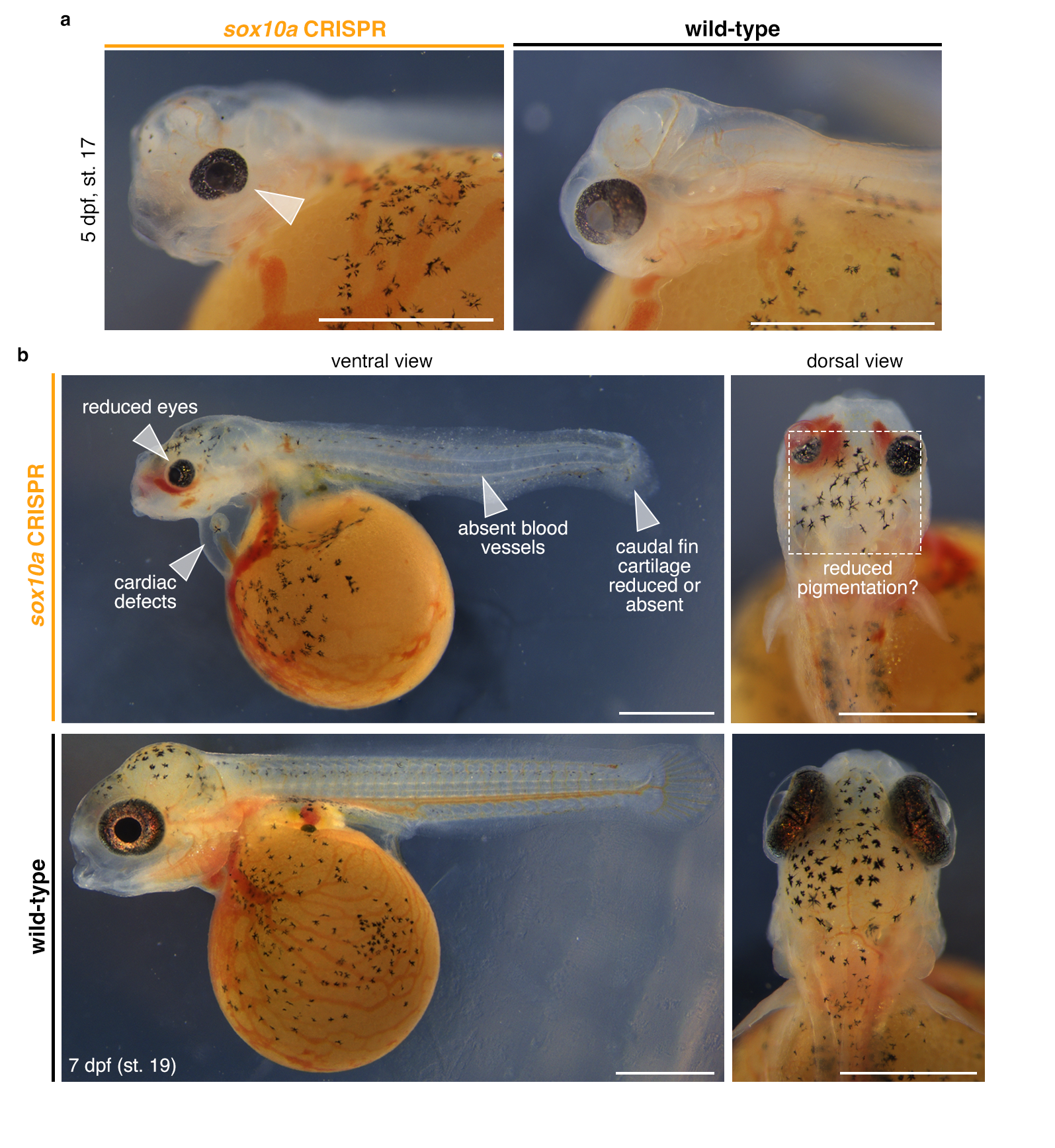

**Supplementary Figure 5. *sox10a* CRISPR embryos display multiple developmental defects. a,** Ocular defects, particularly reduced eye size (grey arrowhead), were observed in *sox10a* KO embryos by 5 dpf (st. 17). **b**, In addition to craniofacial cartilage and ocular malformations, embryos inspected at 7 dpf (st. 19) were also characterised by aberrations of cardiac and circulatory system as well as caudal fin cartilage development. The melanophore-based pigmentation of the dorsal cranium also appeared reduced compared to wild-type, however this could be explained by the reduced surface area of the pigmented region in mutants with severe craniofacial phenotypes. Dpf - days post-fertilisation; st - stage. Scale bars = 1mm.

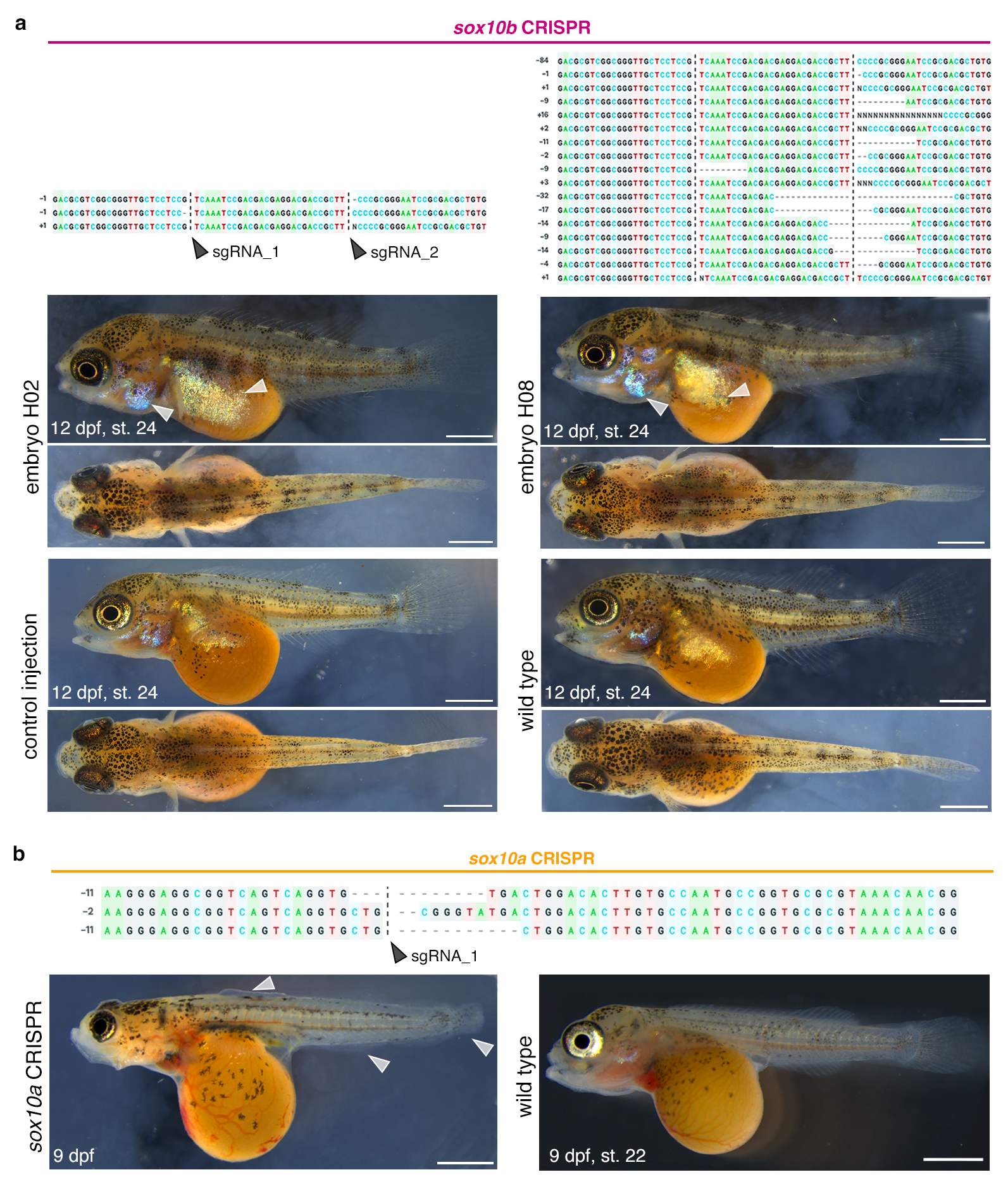

**Supplementary Figure 6.** **Examples of CRISPR/Cas9-induced mutant genotypes and associated phenotypes in *Astatotilapia calliptera* ‘Mbaka’.** Multiple sequence alignments of Sanger sequencing, showing deletions or insertions in exon 1 of *sox10b* (**a**) and *sox10a* (**b**), frequently resulting in frameshifts. **a,** *sox10b* CRISPR embryos at 12 dpf (st. 24, top row) have mild pigmentation abnormalities compared to control and wildtype clutch mates (bottom row), including increased iridophore coverage on the yolk and operculum (grey arrowheads). **b,** In addition to defects of the craniofacial cartilages **(Fig. 6 main text),** surviving *sox10a* CRISPR fish have drastically reduced dorsal, anal and caudal fins (grey arrowheads), likely due to abnormal development of the cartilaginous fin rays.
